# Impact of Realistic 3D Tumor Microarchitecture on Cellular Dosimetry and Radiobiological Response in Targeted Radionuclide Therapy

**DOI:** 10.64898/2026.09.23.753948

**Authors:** Treshan Peirispulle, Robert Ware, Steven Korevaar, Prabath Hetti Mudiyanselage, John Thangarajah, Ruwan Tennakoon

## Abstract

**Background:** Patient-specific dosimetry in targeted radionuclide therapy (TRT) is increasingly supported by quantitative imaging; however, voxel-scale dose estimates cannot resolve cellular and subcellular heterogeneity present within tumor tissue. Conventional cellular dosimetry models often represent cells as regularly packed spheres, which may not capture realistic tumor microarchitecture. This study investigated how histopathology-informed three-dimensional (3D) tumor microarchitecture influences cellular absorbed-dose distributions, predicted radiobiological response, and treatment-related conclusions in TRT.

**Methods:** Realistic 3D cellular tumor models were constructed from thick-section 3D cyclic immunofluorescence (3D CyCIF) imaging data. Monte Carlo radiation-transport simulations were performed for ^177^Lu, ^103^Pd, ^161^Tb and ^90^Y, with activity localized in cancer-cell subcellular compartments and in non-cancer targets. Absorbed dose per total cluster decay was calculated individually for each cell nucleus and compared with corresponding simplified spherical tumor models. Cell survival fractions were estimated using the linear-quadratic model, and tumor control probability (TCP) was calculated from the individual cancer-cell survival probabilities.

**Results:** Simplified model overestimated mean cancer-cell nuclear absorbed dose by **9.54%** to **14.51%** for nuclear localization and by **33.23%** to **44.88%** for cytoplasmic and membrane localization. In larger tissue models, simplified and realistic geometries produced comparable mean absorbed doses in several cases; however, the realistic models consistently generated broader cellular dose distributions, including low-dose cancer-cell subpopulations not captured by the simplified models. These dose-distribution differences propagated into wider survival-fraction distributions and altered TCP predictions. In the spatially heterogeneous tumor region, the realistic model required **1.11**- to **1.33**-fold higher activity to reach ***TCP*_90_** for nuclear localization, and **1.49**- to **5.93**-fold higher activity for cytoplasmic and membrane localization. For similar tumor control, the radionuclide and source-localization combination associated with the greatest preservation of surrounding non-cancer cells differed between the simplified and realistic models. The intra-voxel analysis further showed that homogeneous voxel-scale activity assumptions can obscure biologically relevant microscopic dose heterogeneity.

**Conclusion:** Native 3D tumor microarchitecture substantially influences cellular absorbed-dose distributions, predicted biological response, and treatment-related conclusions. Histopathology-informed 3D cellular models can be used as a complementary framework for characterizing cellular dose heterogeneity and may improve the evaluation of radionuclides and subcellular targeting strategies in heterogeneous tumor tissue.

## 1 Background

Targeted radionuclide therapy (TRT) delivers cytotoxic radiation to malignant tissue through radiopharmaceuticals that localize in tumor-associated molecular targets, enabling irradiation at cellular and subcellular scales while sparing normal tissue [1]. The targeting vehicle determines where the radiopharmaceutical accumulates, whereas the emission spectrum and decay properties of the radionuclide govern the spatial range and magnitude of energy deposition within the tissue [2–5]. The clinical success of TRT is well established: in the phase 3 VISION trial, adding [^177^Lu]Lu – PSMA – 617 to standard of care extended median overall survival from 11.3 to 15.3 months and reduced the risk of death by 38% in patients with metastatic castration-resistant prostate cancer who had exhausted chemotherapy and androgen-receptor-pathway inhibitors [6].

Absorbed dose is a key determinant of both therapeutic response and normal-tissue toxicity in TRT [7]. Effective treatment therefore requires that the administered activity deliver a therapeutically sufficient dose to malignant tissue while limiting irradiation of surrounding normal tissues. Clinically, quantitative SPECT and PET imaging provide patient-specific, macroscopic estimates of the distribution of radio-pharmaceutical activity in tumors and critical organs. Absorbed dose distributions can then be calculated from these activity maps using voxel S-values, dose-point kernel convolution, or Monte Carlo radiation transport methods [2, 8]. However, the spatial resolution of quantitative imaging remains insufficient to capture the cellular energy depositions, leaving microscopic dose heterogeneity arising from variations in cell type, cell packing, and subcellular activity localization unresolved [9]. Consequently, voxel-based dosimetry commonly treats each voxel as a homogeneous volume with a uniform activity concentration and material composition. Characterizing dose deposition beyond the resolution of the imaging voxel is therefore critical for understanding how these sub-voxel variations influence biological response in TRT.

At the cellular scale, in-silico studies evaluating absorbed dose to radiosensitive targets such as the nucleus, support the assessment of promising radionuclides and targeting strategies before experimental or clinical validation [4, 5, 10, 11]. The Medical Internal Radiation Dose (MIRD) schema provides a general framework for radionuclide dosimetry through standardized S-values, which quantify the mean absorbed dose to a target region per radioactive disintegration in a source region, with tools such as MIRDcell extending this framework to multicellular dosimetry and bioeffect modeling across different radionuclides and source-target configurations [12, 13]. Traditionally, these approaches rely on simplified geometrical models, representing individual cells as concentric spheres with a spherical nucleus and approximating tumor clusters using uniform cell size, regular packing, and idealized morphology (Fig. 1b,d). Such spherical geometries may reasonably approximate spheroid cultures, but native tumor tissue can exhibit far greater intratumoral and interpatient heterogeneity [14].

**Fig. 1:**
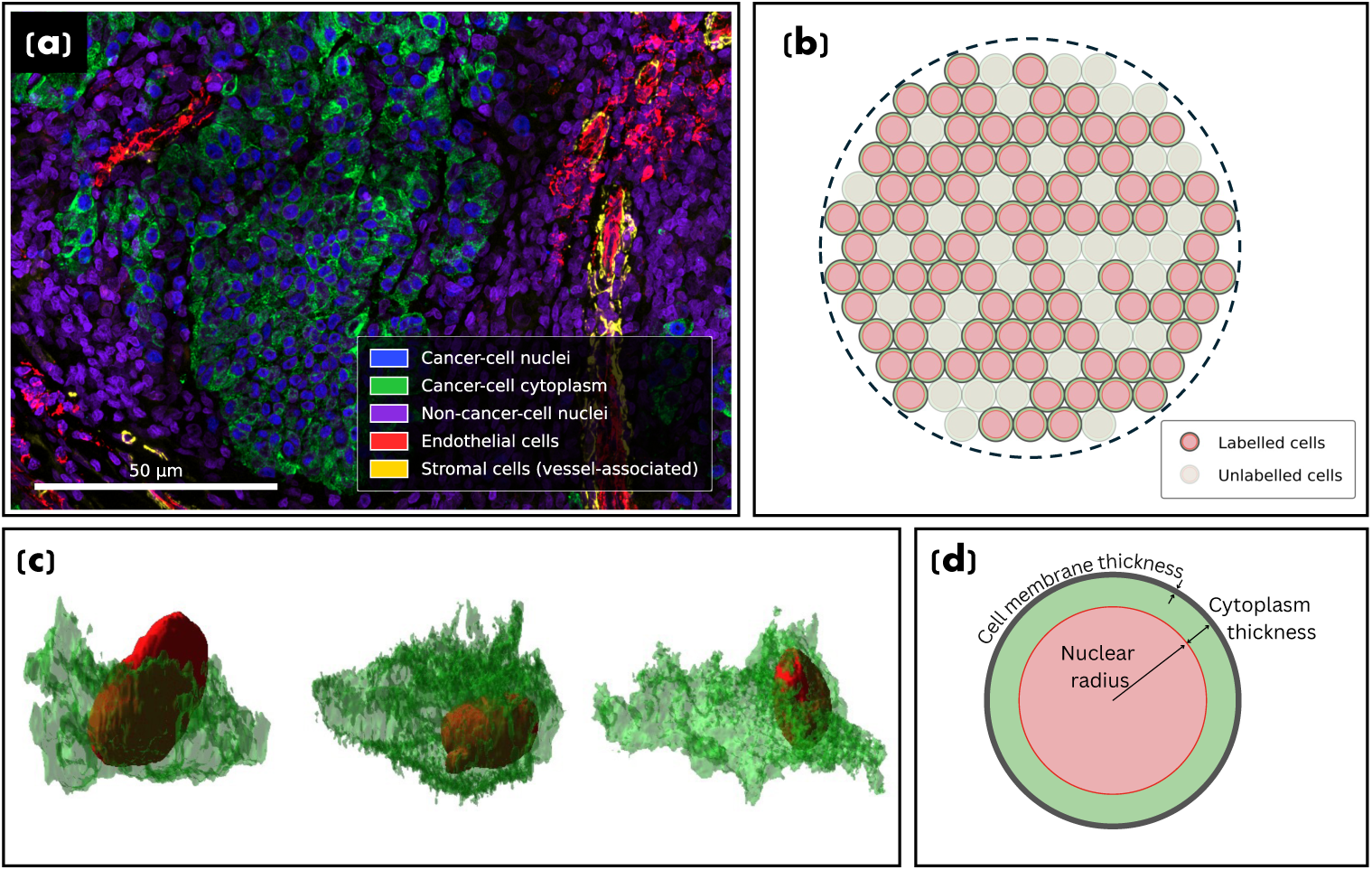
Simplified versus histopathology-informed cellular geometries (a) Maximum-intensity projection of the 3D CyCIF tissue volume. (b) Cross section of simplified multicellular cluster model in which cells are represented as tightly packed, uniformly sized spheres, with a subset labeled with activity. (c) Image-derived realistic cellular morphologies segmented from the CyCIF volume, showing the irregular nucleus (red) and cytoplasm (green) regions. (d) Simplified spherical single-cell geometry.

Cellular dosimetry studies incorporating realistic cancer-cell geometries have shown that absorbed dose estimates are sensitive to modeling assumptions, including cell shape, size, subcellular source localization, clustering, and tumor morphology [10, 11, 15, 16]. These studies consistently report dose distributions that differ from those predicted by spherical MIRDcell-type geometries, with recent work suggesting that cell clustering and tissue scale morphology may have a greater influence on absorbed dose estimates than isolated single cell geometry [11, 16]. However, most existing models are derived from isolated cells, two-dimensional tumor maps, or in vitro cultures, limiting their ability to capture realistic three-dimensional (3D) cross-fire effects and heterogeneous cellular compositions within native tumor microenvironments [10, 11, 16]. Consequently, how the native 3D tumor microenvironment alters cellular absorbed dose estimates and downstream radiobiological response predictions in TRT remains insufficiently characterized.

Recent advances in high-resolution optical microscopy and multiplexed tissue imaging offer an opportunity to address these limitations. In particular, thick-section 3D cyclic immunofluorescence (3D CyCIF) enables volumetric imaging of intact tumors while simultaneously profiling dozens of molecular markers, allowing cancer and non-cancer cell populations to be identified within a preserved tissue environment [14]. This provides a more realistic view of the tumor microenvironment, distinguishing cancer cells from the diverse non-cancer populations that surround them, including immune cells, endothelial cells, and cancer-associated stromal cells (Fig. 1a). This makes 3D CyCIF well-suited for constructing realistic cellular and subcellular dosimetry models that incorporate native tumor architecture and heterogeneous cellular composition.

In this work, we develop realistic 3D computational tumor phantoms from CyCIF data to investigate how native tumor microarchitecture influences cellular-scale dosimetry and radiobiological response in TRT. To our knowledge, this is the first study to translate thick-section CyCIF imaging into 3D cellular dosimetry models. Using Monte Carlo simulations, we calculate nucleus absorbed doses for different radionuclides and source localizations and estimate corresponding survival fractions and tumor control probabilities (TCP). By comparing these results with conventional MIRDcell-type geometries, we evaluate whether histopathology-informed tumor models produce meaningful differences in absorbed dose distributions and treatment-response predictions. This framework provides a foundation for incorporating realistic tissue architecture into cellular-scale TRT dosimetry.

## 2 Methods

### 2.1 3D CyCIF Data

A publicly available 3D CyCIF dataset by Yapp et al. [14] was used to construct the realistic tumor models. In the original study, a 35 *µ*m-thick tissue section from an invasive vertical growth phase (VGP) melanoma specimen was imaged using high-resolution confocal microscopy, resulting in a 54-plex 3D CyCIF volume. Thick-section imaging enabled visualization of intact cancer, immune, and stromal cells, along with their native local spatial organization. The dataset was available as a multiresolution image pyramid, from which the highest-resolution representation, with voxel spacing of 0.28 *µ*m, 0.14 *µ*m, and 0.14 *µ*m in the *Z*, *Y*, and *X* directions respectively, was selected for model generation.

### 2.2 Image Preprocessing and Cellular Segmentation

Image preprocessing followed the same procedure described by Yapp et al. [14]. Uneven illumination was corrected by estimating a smooth background from a downsampled and Gaussian-filtered version of the 3D image volume, which was resampled to the original image size and used to normalize the raw image to obtain an illumination-corrected volume. Contrast stretching was applied using the 2nd and 99.8th percentile intensity values, followed by clipping to the range [0, 1], with all steps performed at the native spatial resolution of the data.

Cellpose 3D segmentation model was applied to the Hoechst channel after preprocessing to segment all nuclei [17]. Post-processing was performed to remove implausible segmentations based on object size. Small objects with fewer than 4000 voxels (21.95 *µ*m^3^) were discarded, and unusually large objects were removed using a statistical threshold defined as the mean nuclear volume plus five standard deviations. Only objects satisfying both criteria were retained in the final segmentation mask, excluding noise, fragmented nuclei, and merged or over-segmented objects.

Cancer-cell nuclei were identified based on SOX10 expression on the segmented nuclei, and cytoplasmic regions were delineated using the MART1 channel together with the nuclear masks. Watershed segmentation was applied using the cancer-cell nuclei as seeds and the negative MART1 intensity image as the gradient, separating adjacent cytoplasmic regions. Membrane-associated regions were subsequently defined as the boundary voxels of the segmented cancer-cell regions using six-connectivity in the 3D voxel grid. The resulting nuclear, cytoplasmic, and membrane-associated compartments were used for subsequent model construction and dosimetric analysis (Additional file 1: Fig. S1).

### 2.3 Generation of Simplified Tumor Models

The segmented data were used to estimate the mean cellular diameter (*D_cell_* = 10.38 *±* 2.79 *µ*m) and mean nuclear diameter (*D_nuc_* = 7.95 *±* 1.53 *µ*m), which were used to construct Simplified Models 1, 2, and 3, ensuring that the simplified geometry remained representative of the realistic image-derived phantoms and enabling direct comparison between the two models.

#### Simplified Model 1

was designed to isolate the effect of realistic cell geometry and spatial organization by assuming a homogeneous cluster composed entirely of cancer cells. Following the configuration described by Hindié et al. [4], the model consisted of a central cancer cell surrounded by six first-neighbor cells and twelve second-neighbor cells, totalling 19 cancer cells (Fig. 2a). For subsequent Monte Carlo simulations, activity was assigned uniformly to one of three subcellular compartments: the nucleus, cytoplasm, or membrane-associated region.

**Fig. 2:**
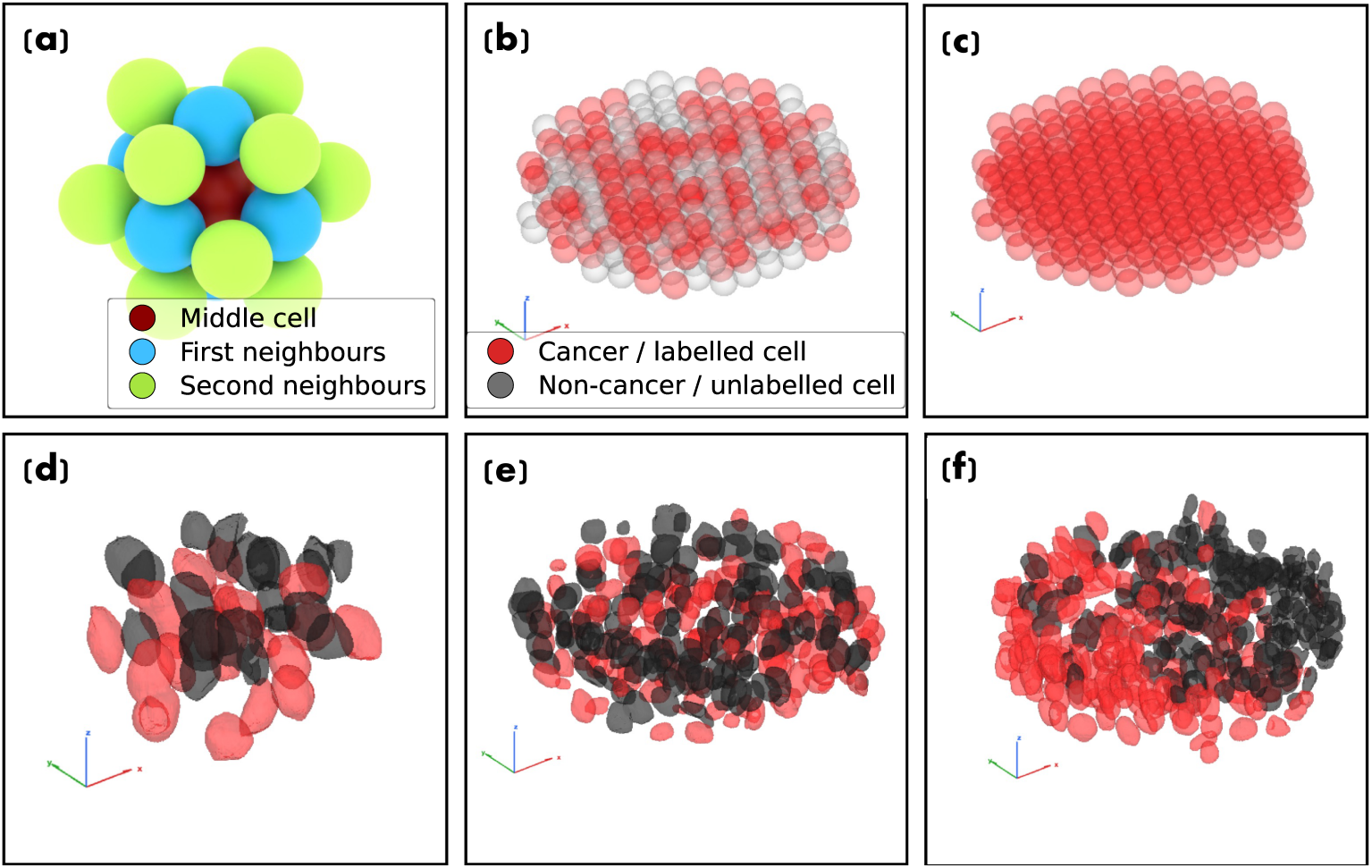
Simplified and image-derived realistic tumor models used for cellular-scale dosimetry. (a) *Simplified Model 1* : a homogeneous 19-cell cluster with a central cancer cell surrounded by six first-neighbor and twelve second-neighbor cells, each represented as a sphere with a concentric spherical nucleus. (b) *Simplified Model 2* : tightly packed spherical cells randomly labeled as cancer (red) or non-cancer (gray) cells, reproducing the cancer-cell fraction of the corresponding realistic model. (c) *Simplified Model 3* : a uniformly labeled cluster in which all cells are designated cancer cells, representing an idealized homogeneous intra-voxel activity distribution. (d–f) Corresponding 3D CyCIF-derived realistic cellular geometries: (d) *Realistic Model 1* : cluster with 19 cancer cells; (e) *Realistic Model 2* : an intermixed population of cancer and non-cancer cells; (f) *Realistic Model 3* : a spatially heterogeneous region in which cancer cells form a concentrated subregion.

#### Simplified Model 2

was developed to account for the heterogeneous cellular composition of the tumor microenvironment. The model consisted of tightly packed spherical cells distributed within an overall cluster volume equal to that of realistic models. The cancer-cell fraction calculated in each realistic model was reproduced by randomly labeling the same proportion of cells as cancer cells, with the remaining cells as non-cancer cells (Fig. 2b).

#### Simplified Model 3

was constructed to examine the implications of an idealized, homogeneous intra-voxel activity distribution. A SPECT/PET voxel with uniform activity distribution can be conceptualized as a tissue volume containing uniformly distributed radiolabeled cancer cells. To approximate this assumption, tightly packed spherical cells were distributed within the same total cluster volume as the realistic models, with all cells labeled as cancer cells (Fig. 2c). This model served as a reference case for examining intra-voxel heterogeneity, through comparison with the realistic tissue-derived models in which cancer cells were heterogeneously distributed and interspersed with non-cancer cells.

### 2.4 Generation of Realistic 3D Tumor Models

Three types of realistic tumor models: Realistic Models 1, 2, and 3, were generated from the segmented 3D CyCIF dataset.

#### Realistic Model 1

was extracted by selecting a tissue volume containing 19 cancer cells (Fig. 2d). This model served as the realistic counterpart to the *Simplified Model 1*. To assess variability across tissue regions and support statistical comparison with the simplified model a total of 20 similar image-derived clusters were generated using the same procedure.

Two larger rod-shaped realistic models were also generated from selected regions of the segmented 3D CyCIF dataset, each with a radius of 61 *µ*m and a height of 30 *µ*m.

#### Realistic Model 2

contained a relatively intermixed population of cancer and non-cancer cells distributed throughout the selected volume (Fig. 2e).

#### Realistic Model 3

represented a more spatially heterogeneous tissue region, in which cancer cells formed a concentrated subregion and non-cancer cells occupied a larger proportion of the surrounding volume (Fig. 2f). Together, these models were used to investigate how differences in cellular composition and spatial organization influence absorbed-dose distributions and predicted radiobiological response. A comparison of the geometrical and cellular properties of the simplified and image-derived realistic models is provided in the Additional file 1: Table S1.

*Realistic Model 3* was additionally used to examine cross-irradiation effects arising from non-specific radiopharmaceutical uptake. For this analysis, activity was localized within the non-cancer-cell population rather than the cancer-cell compartments, and the resulting absorbed-dose distributions to both cancer-cell and non-cancer-cell nuclei, along with the corresponding predicted radiobiological response, were evaluated and compared with the cancer-cell-targeted source localizations.

### 2.5 Monte Carlo Simulation of Cellular Absorbed Dose

The Monte Carlo simulation software GATE (v 10.0), based on Geant4 (v 11.4.0), was used to simulate radioactive decay and particle transport [18]. Four radionuclides were modeled: relatively long-range *β^−^* emitting ^177^Lu and ^90^Y, low-energy Auger-electron emitting ^103^Pd, and ^161^Tb, which supplements its *β^−^* decay with substantial short-range conversion and Auger electrons. The decay properties of each radionuclide are summarized in Additional file 1: Table S9. For each model simulations were performed separately for each radionuclide, with activity distributed uniformly within the corresponding compartment. Particle transport was modeled using the G4EmLivermorePhysics electromagnetic physics list. Radioactive decay was enabled, with daughter radionuclides included in the simulations. Atomic de-excitation processes, including fluorescence emission, Auger-electron emission, and Auger cascades, were enabled to account for short-range electron emissions. Production cuts of 0.01 *µ*m were applied to electrons, and photons. For each combination of model, radionuclide, and source localization, 10^6^ radionuclide decays were simulated. The mean percentage dose uncertainty per nucleus for each radionuclide and source localization is provided in the Additional file 1: Table S2. The absorbed dose delivered to each cell nucleus was recorded separately for further analysis.

### 2.6 Cell Survival and Tumor Control Probability Modeling

The Monte Carlo absorbed dose results were normalized and expressed as the mean nuclear absorbed dose per total cluster decay. For cell *i*, this quantity was denoted by *d_i_* (Gy decay*^−^*^1^). Unless stated otherwise, all absorbed dose values reported hereafter refer to the mean nuclear absorbed dose normalized per total cluster decay. To estimate the cumulative absorbed dose for a given initial cluster activity, the total number of radionuclide decays was calculated using the radionuclide-specific time-integrated activity coefficient (TIAC), assuming no biological clearance from the tumor. For radionuclide *r*, the total number of decays was calculated as

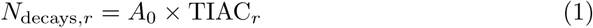

where *A*_0_ is the initial cluster activity in Bq and TIAC*_r_* is the time-integrated activity coefficient of radionuclide *r*, converted to seconds. The TIAC values and the corresponding total cluster decays for *A*_0_ = 0.08 Bq and *A*_0_ = 0.1 Bq are listed in Table 1. The cumulative absorbed dose to the nucleus of cell *i* was then calculated as

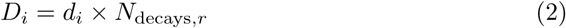

where *D_i_* is expressed in Gy.

**Table 1:** Time-integrated activity coefficients and corresponding total cluster decays for selected initial cluster activities, assuming no biological washout.

| Radionuclide | TIAC (h) | Total cluster decays<br>($A_0 = 0.08$ Bq) | Total cluster decays<br>($A_0 = 0.1$ Bq) |
| --- | --- | --- | --- |
| $^{177}\text{Lu}$ | 230.20 | $6.63 \times 10^4$ | $8.29 \times 10^4$ |
| $^{103}\text{Pd}$ | 588.43 | $1.69 \times 10^5$ | $2.12 \times 10^5$ |
| $^{161}\text{Tb}$ | 239.12 | $6.89 \times 10^4$ | $8.61 \times 10^4$ |
| $^{90}\text{Y}$ | 92.50 | $2.66 \times 10^4$ | $3.33 \times 10^4$ |

The cumulative absorbed dose to each cell nucleus was used to estimate the corresponding cell survival fraction using the linear-quadratic model. For a cell receiving a mean absorbed dose *D_i_*, the survival fraction was calculated as

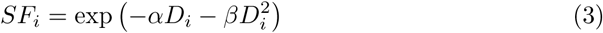

where *SF_i_* is the survival fraction of cell *i*, and *α* and *β* are the radiosensitivity parameters. For cancer cells, *α* = 0.37 Gy*^−^*^1^ and *β* = 0.06 Gy*^−^*^2^ were used based on previously reported clonogenic survival data for a human primary melanoma cell line [19]. For non-cancer cells, reference values of *α* = 0.30 Gy*^−^*^1^ and *β*= 0.03 Gy*^−^*^2^ were used to enable comparative assessment of irradiation between tumor models. The survival fraction, *SF_i_*, represents the probability that cancer cell *i* remains capable of regrowth following irradiation. Survival fraction was calculated separately for each cancer cell, characterizing the influence of cellular-scale dose heterogeneity on the predicted radiobiological response.

Although the survival fraction provides a measure of cell-level response, it does not fully describe tumor control, which depends on the number of surviving cancer cells and their potential to repopulate the tumor [20]. TCP was therefore estimated from these survival fractions, assuming independent cell survival, as

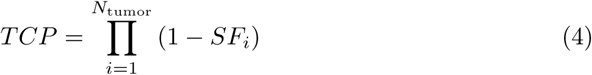

where *N*_tumor_ is the number of cancer cells in the modeled cluster.

## 3 Results

### 3.1 Simplified Tumor-Cluster Models Overestimate Nuclear Absorbed Dose

Mean cancer-cell absorbed dose for all radionuclides and source localizations was calculated separately for each of the 20 independently sampled *Realistic Model 1* clusters and compared with the *Simplified Model 1* (Fig. 3a ; Additional file 1: Table S3). Across all radionuclides and source localizations, ^161^Tb produced the highest mean nuclear dose, followed by ^103^Pd, ^177^Lu and ^90^Y. For all radionuclide and source-localization combinations, the simplified-cluster mean absorbed dose consistently exceeded the realistic-model value; the associated two-sided one-sample *t*-test *p*-values were below 0.05 in each case (Additional file 1: Table S3). The magnitude of the difference depended on the subcellular source localization. Realistic clusters received *∼*10–16% lower mean doses for nuclear localization, but 33–45% lower doses for cytoplasmic and membrane localization, indicating that the simplified geometry overestimates dose the most when activity resides outside the nucleus.

**Fig. 3:**
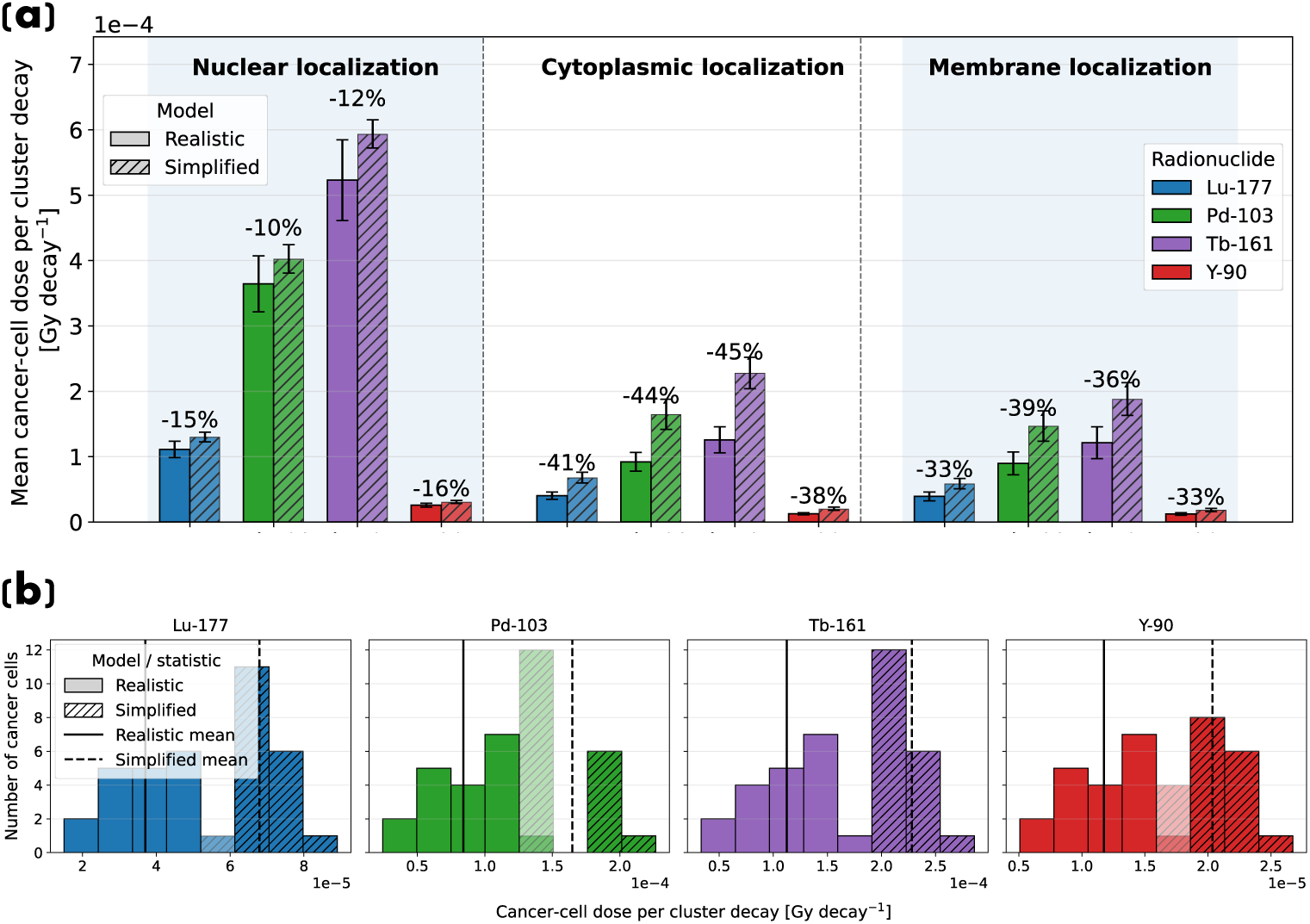
Comparison of cancer-cell nuclear absorbed dose between *Simplified Model 1* and *Realistic Model 1*. (a) Mean absorbed dose to cancer-cell nuclei per cluster decay. Error bars indicate the standard deviation across the 20 sampled realistic clusters. Percentage labels denote the relative difference in mean dose for the realistic model compared with the simplified model. (b) Cancer-cell nuclear absorbed-dose distributions for cytoplasmic source localization. Semi-transparent bars denote overlapping dose bins between the two distributions.

Cancer-cell nuclear dose distribution histograms for cytoplasmic source localization are shown in Fig. 3b. Because *Simplified Model 1* has a regular radially symmetric arrangement, its dose distribution is concentrated in a narrow dose range associated with the central cell, first-neighbor cells, and second-neighbor cells. In contrast, realistic clusters lack this regular spatial organization and show broader dose distributions arising from variations in cell morphology and local cellular arrangement.

### 3.2 Impact of Tumor Microarchitecture on Nuclear Absorbed Doses

A consistent pattern emerged across large cluster comparisons: mean nuclear absorbed doses were often comparable between the simplified and realistic models, but the realistic geometry produced substantially broader cell-to-cell dose distributions in every case. This effect was most pronounced for ^103^Pd and ^161^Tb, consistent with their shorter-range emissions, which increase sensitivity to local variations in cell morphology and spacing.

#### 3.2.1 Spatially homogeneous tumor region (*Realistic Model 2*)

Mean absorbed doses to cancer and non-cancer cell nuclei were compared between *Simplified Model 2* and *Realistic Model 2* (Fig. 4a-b; Additional file 1: Table S4). For cancer cells, the effect of realistic microarchitecture depended on subcellular source localization: the realistic model produced higher mean nuclear doses for nuclear localization (+14% to +27%), but lower doses for cytoplasmic (*−*16% to *−*24%) and membrane (*−*9% to *−*12%) localization (Fig. 4a). Mean non-cancer-cell doses differed only modestly between models (*−*15% to +17%; Fig. 4b). In both populations, however, the realistic model produced markedly broader dose distributions across individual cells, consistent with the general pattern noted above. Matching the cancer-cell fraction in a regularly packed model can therefore approximate the mean absorbed dose, but does not reproduce the cellular dose heterogeneity arising from realistic morphology and spatial organization, a distinction relevant for tumor-control analysis, which depends on the survival of individual cells.

**Fig. 4:**
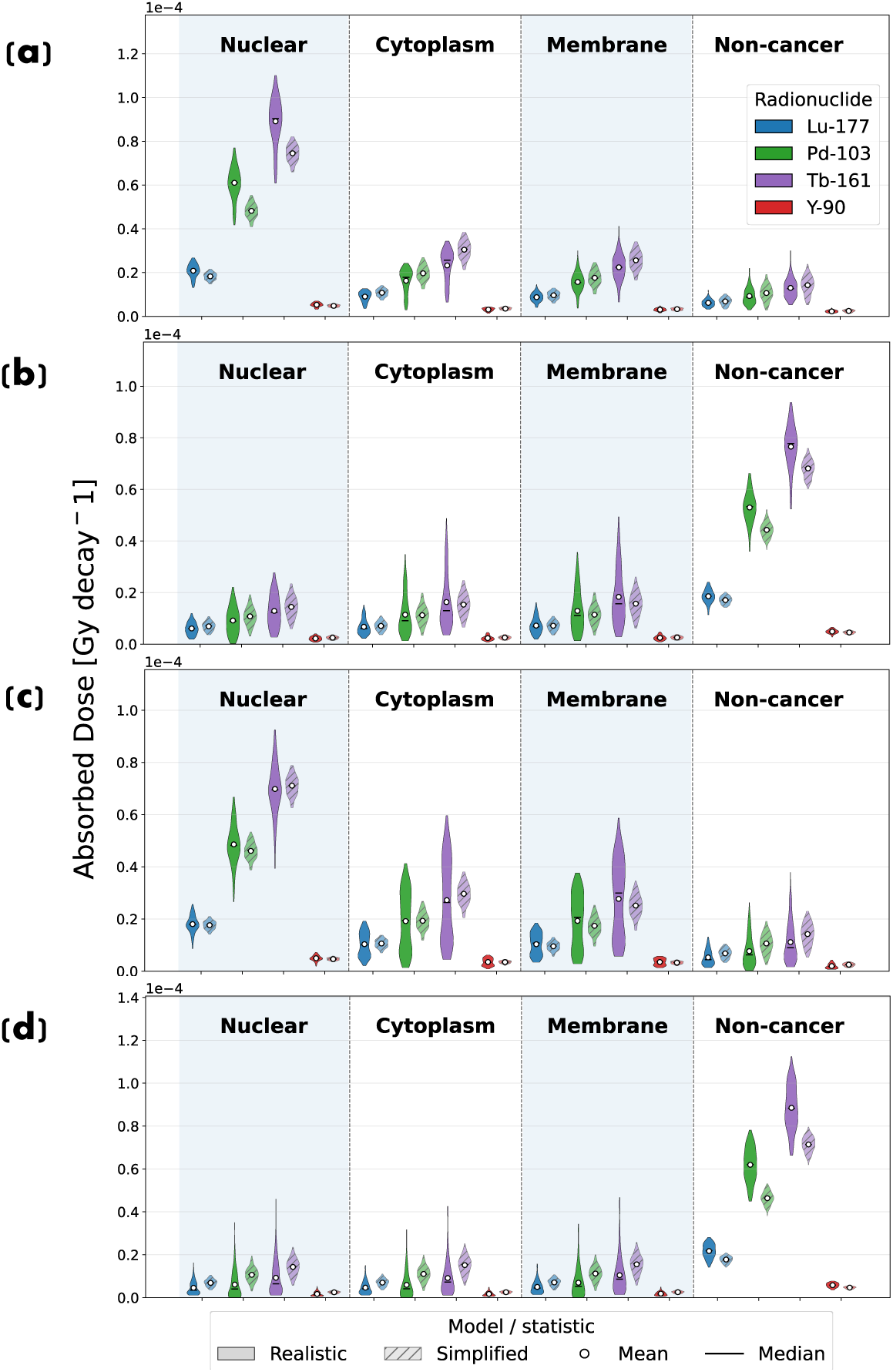
Distributions of nuclear absorbed dose per cluster decay for cancer-cell and non-cancer-cell nuclei, comparing *Simplified Model 2* with the image-derived realistic tumor models. (a) Cancer-cell nuclear dose distributions for *Simplified Model 2* and *Realistic Model 2*. (b) Non-cancer-cell nuclear dose distributions for the same model pair. (c) Cancer-cell nuclear dose distributions for *Simplified Model 2* and *Realistic Model 3*. (d) Non-cancer-cell nuclear dose distributions for the same model pair. Column headers Nuclear, Cytoplasm, Membrane, and Non-cancer denote activity localized in the cancer-cell nucleus, cytoplasm, and membrane-associated compartment, and in non-cancer cells, respectively.

#### 3.2.2 Spatially heterogeneous tumor region (*Realistic Model 3*)

In *Realistic Model 3*, cancer cells were not uniformly intermixed with non-cancer cells but instead formed a more concentrated subregion. Mean cancer-cell nuclear doses were again broadly comparable to the randomly labeled simplified model, with differences ranging from an 8% reduction to an 11% increase in subcellular localizations (Fig. 4c; Additional file 1: Table S5a), while the cellular dose distributions were substantially broader in the realistic geometry revealing both low-dose and high-dose cancer-cell subpopulations not captured by the simplified model. In contrast, mean doses to noncancer-cells were consistently lower in the realistic model, except when radioactivity was localized in non-cancer cells, with reductions of 34–47% (Fig. 4d; Additional file 1: Table S5b), and again showed substantial cell-to-cell variation.

As with the on-target cancer-cell doses, the cross-dose to tumor-cell nuclei was highest for ^161^Tb, followed by ^103^Pd, ^177^Lu and ^90^Y when activity is localized in non-cancer cells. Compared with directly targeting cancer cells, activity localized in non-cancer cells delivered consistently lower doses to cancer-cell nuclei. In realistic model this cross-dose corresponded to 29.2%, 50.9%, and 51.0% of the dose achieved by direct nuclear, cytoplasmic, and membrane localization respectively, for ^177^Lu; 15.8%, 40.0%, and 39.8% for ^103^Pd; and 15.9%, 40.9%, and 40.2% for ^161^Tb.

### 3.3 Cellular Dose Heterogeneity Influences Predicted Radiobiological Response

The probability that a cell survives irradiation, expressed as its survival fraction, and the tumor control probability (TCP) were evaluated at different initial cluster activities. Across radionuclides, ^103^Pd consistently produced the lowest mean survival probability, followed by ^161^Tb, ^177^Lu, and ^90^Y. The absorbed-dose patterns identified in §3.2 propagated directly into the radiobiological endpoints: differences in mean nuclear dose were reflected in mean survival fractions, while the broader cellular dose distributions produced broader survival-fraction distributions in the realistic models.

#### 3.3.1 Survival fractions (*Realistic Model 2*)

Cancer-cell survival fractions for *Realistic Model 2* and *Simplified Model 2* at an initial cluster activity of *A*_0_ = 0.08 Bq are shown in Fig. 5a (Additional file 1: Table S6a). For cytoplasmic, membrane, and non-cancer-cell localizations, cancer cells in the realistic model were more likely to survive irradiation than in the simplified model when comparing mean values, consistent with the lower mean nuclear doses in those configurations (Fig. 4a). However, the corresponding survival-fraction distributions revealed subpopulations of cells with both a higher probability of being killed and a higher probability of survival than predicted by the simplified model.

**Fig. 5:**
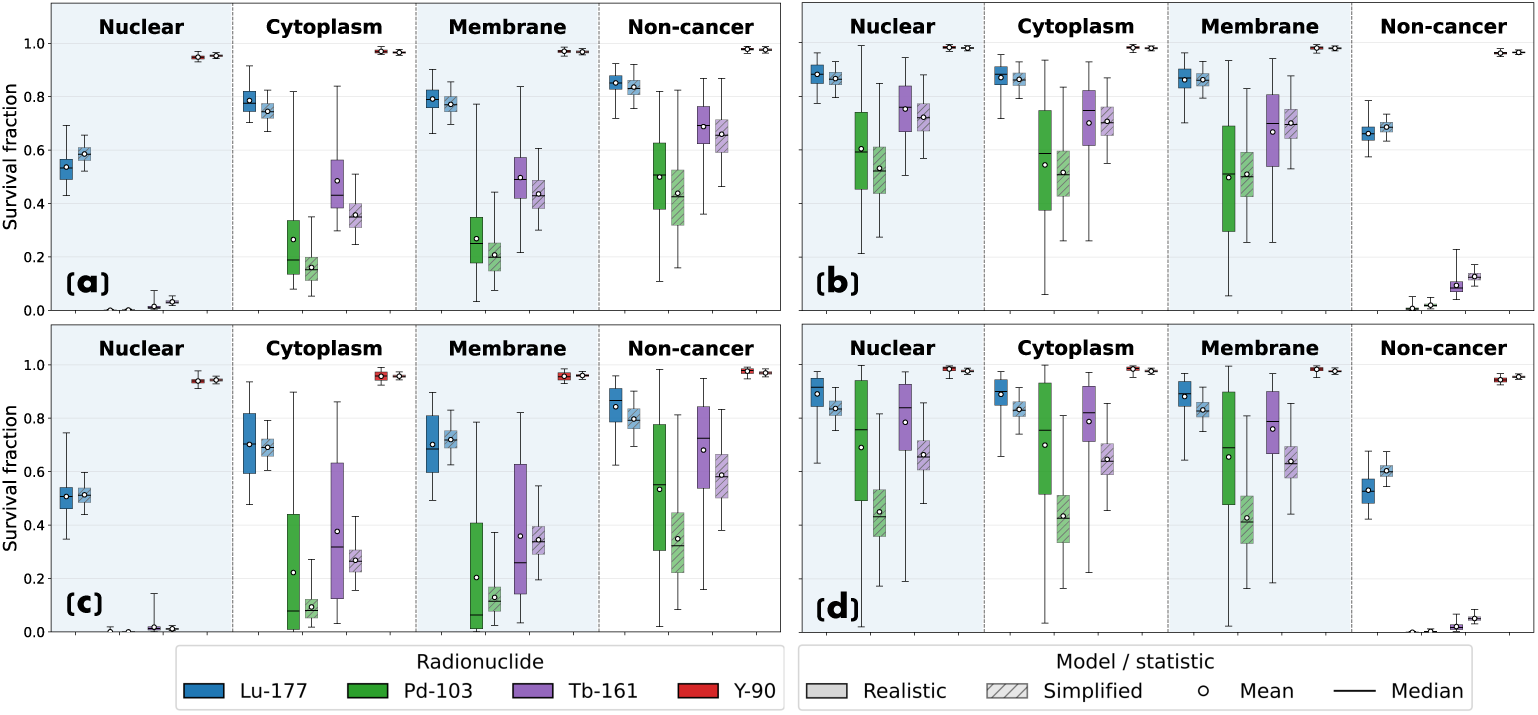
Comparison of predicted cellular survival fractions (a) Cancer-cell survival fractions, (b) Non-cancer cell survival fractions in the *Simplified Model 2* and image-derived *Realistic Model 2* at an initial total cluster activity of *A*_0_ = 0.08 Bq. (c) Cancer-cell survival fractions, (d) Non-cancer cell survival fractions in the *Simplified Model 2* and image-derived *Realistic Model 3* at an initial total cluster activity of *A*_0_ = 0.1 Bq. Column headers Nuclear, Cytoplasm, Membrane, and Non-cancer denote activity localized in the cancer-cell nucleus, cytoplasm, and membrane-associated compartment, and in non-cancer cells, respectively.

Mean non-cancer-cell survival fractions were comparable between models, but again with broader distributions in the realistic geometry (Fig. 5b; Additional file 1: Table S6b), particularly for ^103^Pd and ^161^Tb.

#### 3.3.2 Survival fractions (*Realistic Model 3*)

The radiobiological response was next evaluated in the more spatially heterogeneous *Realistic Model 3*, compared with the randomly labeled *Simplified Model 2* at *A*_0_ = 0.1 Bq (Fig. 5c and Fig. 5d; Additional file 1: Table S7). For nuclear localization, both models predicted near-complete cancer-cell killing for ^103^Pd and ^161^Tb, with a more modest difference for ^177^Lu. Cytoplasmic and membrane localization produced larger discrepancies, with the realistic model predicting higher probability of mean survival for cancer cells. The resulting low- and high-dose cancer-cell subpopulations translate directly into subpopulations of cells that are, respectively, largely spared or almost completely killed, a response not captured by the simplified model’s narrower dose distribution. Non-cancer cells showed the same pattern: higher mean survival fractions in the realistic model, with broader distributions across individual cells (Fig. 5d). Consistent with the lower cross-dose, localizing activity in non-cancer cells produced higher cancer-cell survival fractions than direct cancer-cell targeting for all radionuclides, while resulting in greater killing of the non-cancer cells themselves.

#### 3.3.3 Activity required for *T CP*_90_ and radionuclide ranking

TCP was calculated over a range of initial cluster activities (0.001–20 Bq), and the activity required to reach *TCP*_90_ was determined for each radionuclide and source localization (Table 2). The realistic model required higher activities than the simplified model in every case. Among sub-cellular localizations the smallest differences were observed for nuclear localization (realistic model required 1.11 *−* 1.33*×* the activity of the simplified model), while cytoplasmic and membrane localization required 1.49 *−* 5.93*×* higher activities, the maximum occurring for ^103^Pd with cytoplasmic localization. These increases are consistent with the low-dose cancer-cell subpopulations identified in the corresponding dose distributions (Fig. 4c).

**Table 2:** Initial total cluster activity required to achieve a tumor control probability of 90% (*TCP*_90_) in the *Simplified Model 2* and *Realistic Model 3* tumor models. The ratio is calculated relative to the simplified model.

| Radionuclide | Source localization | Simplified model<br>$A_0$ for $TCP_{90}$ (Bq) | Realistic model<br>$A_0$ for $TCP_{90}$ (Bq) | Realistic /<br>simplified<br>ratio |
| --- | --- | --- | --- | --- |
| $^{177}\text{Lu}$ | Nucleus | 0.601 | 0.743 | 1.24 |
| $^{103}\text{Pd}$ | Nucleus | 0.089 | 0.098 | 1.11 |
| $^{161}\text{Tb}$ | Nucleus | 0.140 | 0.164 | 1.17 |
| $^{90}\text{Y}$ | Nucleus | 5.863 | 7.810 | 1.33 |
| $^{177}\text{Lu}$ | Cytoplasm | 1.094 | 2.449 | 2.24 |
| $^{103}\text{Pd}$ | Cytoplasm | 0.246 | 1.459 | 5.93 |
| $^{161}\text{Tb}$ | Cytoplasm | 0.365 | 1.321 | 3.61 |
| $^{90}\text{Y}$ | Cytoplasm | 8.411 | 15.791 | 1.88 |
| $^{177}\text{Lu}$ | Membrane | 1.266 | 2.079 | 1.64 |
| $^{103}\text{Pd}$ | Membrane | 0.289 | 0.875 | 3.03 |
| $^{161}\text{Tb}$ | Membrane | 0.455 | 1.119 | 2.46 |
| $^{90}\text{Y}$ | Membrane | 9.148 | 13.613 | 1.49 |
| $^{177}\text{Lu}$ | Non-cancer cell | 2.134 | 4.809 | 2.25 |
| $^{103}\text{Pd}$ | Non-cancer cell | 0.891 | 10.113 | 11.36 |
| $^{161}\text{Tb}$ | Non-cancer cell | 1.163 | 3.662 | 3.15 |
| $^{90}\text{Y}$ | Non-cancer cell | 14.023 | —<sup>†</sup> | —<sup>†</sup> |

<sup>†</sup>Within the tested activity range (up to 20 Bq), $TCP_{90}$ was not reached.

In the realistic model, within on-target (cancer-cell) localizations, the required activity followed a consistent radionuclide ordering for nuclear and membrane localization, increasing as ^103^Pd *<* ^161^Tb *<* ^177^Lu *<* ^90^Y; ^90^Y required the greatest activity of the four radionuclides at every source localization, consistent with its comparatively low dose deposition seen earlier (Fig. 4c). For a given radionuclide, membrane localization consistently required lower activity than cytoplasmic localization to reach *TCP*_90_(^177^Lu: 2.079 vs. 2.449 Bq; ^103^Pd: 0.875 vs. 1.459 Bq; ^161^Tb: 1.119 vs. 1.321 Bq), indicating that membrane-localized activity was modestly more effective at achieving tumor control than cytoplasmic localization for all three radionuclides.

Across all radionuclides, non-cancer-cell localization required the highest activity of any source localization, since this scenario depends on cross-dose delivered from neighboring non-cancer cells rather than direct on-target irradiation, resulting in lower absorbed dose to cancer-cell nuclei than any of the cancer-cell-targeted localizations. Radionuclide and source-localization combinations were then ranked by predicted non-cancer-cell kill fraction at *TCP*_90_ (Table 3). Nuclear localization ranked highest in both models, but the preferred radionuclide differed: in the simplified model, ^161^Tb produced the lowest non-cancer kill (44.86%), whereas in the realistic model ^103^Pd ranked first (30.65%). This re-ordering indicates that utilising realistic tumor microarchitecture in simulations alter both the magnitude of predicted outcomes as well as the impact on both cancer and non-cancer cell survival rates, thus likely impacting the final choice of radionuclide for treatment.

**Table 3:**
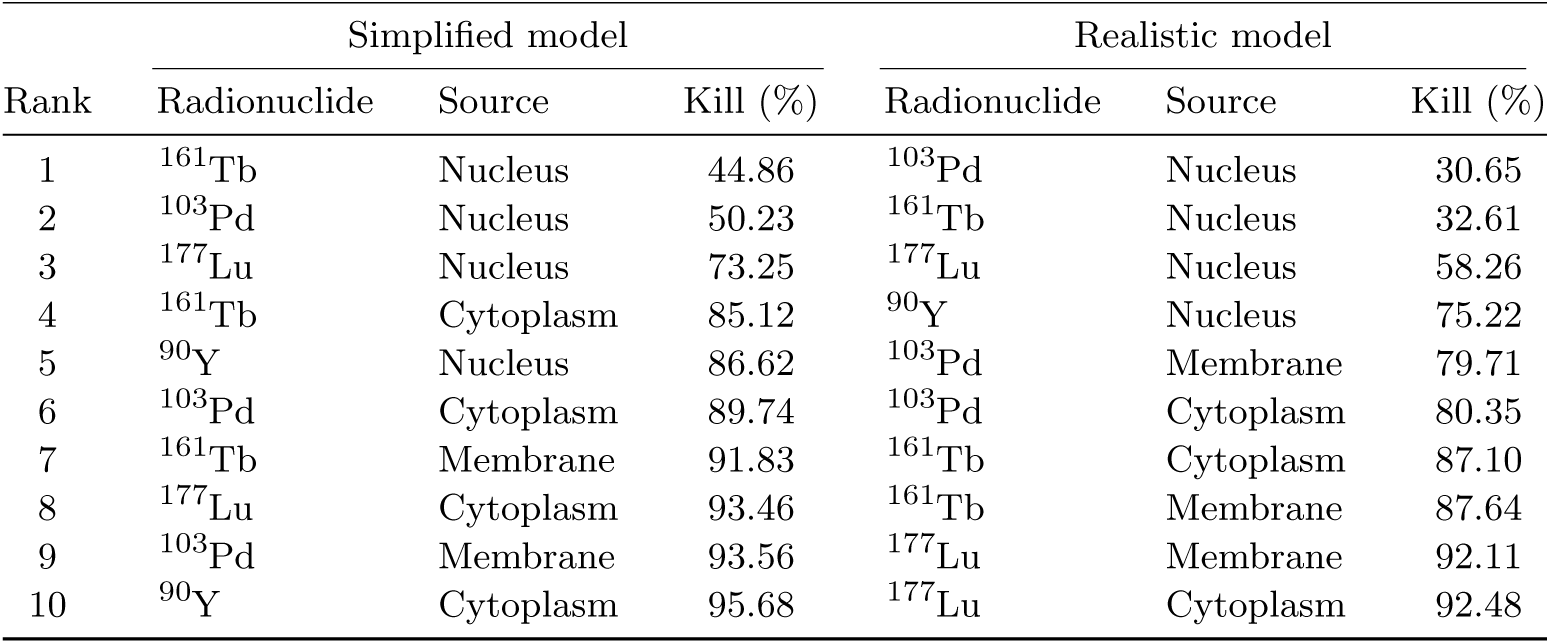
Ranking of radionuclide and source-localization combinations in ascending order of the predicted non-cancer-cell kill fraction at *TCP*_90_ compared between *Simplified Model 2* and *Realistic Model 3*. Lower values indicate greater preservation of surrounding non-cancer cells.

| Rank | Simplified model |  |  | Realistic model |  |  |
| --- | --- | --- | --- | --- | --- | --- |
|  | Radionuclide | Source | Kill (%) | Radionuclide | Source | Kill (%) |
| 1 | $^{161}\text{Tb}$ | Nucleus | 44.86 | $^{103}\text{Pd}$ | Nucleus | 30.65 |
| 2 | $^{103}\text{Pd}$ | Nucleus | 50.23 | $^{161}\text{Tb}$ | Nucleus | 32.61 |
| 3 | $^{177}\text{Lu}$ | Nucleus | 73.25 | $^{177}\text{Lu}$ | Nucleus | 58.26 |
| 4 | $^{161}\text{Tb}$ | Cytoplasm | 85.12 | $^{90}\text{Y}$ | Nucleus | 75.22 |
| 5 | $^{90}\text{Y}$ | Nucleus | 86.62 | $^{103}\text{Pd}$ | Membrane | 79.71 |
| 6 | $^{103}\text{Pd}$ | Cytoplasm | 89.74 | $^{103}\text{Pd}$ | Cytoplasm | 80.35 |
| 7 | $^{161}\text{Tb}$ | Membrane | 91.83 | $^{161}\text{Tb}$ | Cytoplasm | 87.10 |
| 8 | $^{177}\text{Lu}$ | Cytoplasm | 93.46 | $^{161}\text{Tb}$ | Membrane | 87.64 |
| 9 | $^{103}\text{Pd}$ | Membrane | 93.56 | $^{177}\text{Lu}$ | Membrane | 92.11 |
| 10 | $^{90}\text{Y}$ | Cytoplasm | 95.68 | $^{177}\text{Lu}$ | Cytoplasm | 92.48 |

### 3.4 Implications for intra-voxel dose heterogeneity

To assess the implications of intra-voxel heterogeneity, the uniformly labeled *Simplified Model 3* was compared with the corresponding image-derived *Realistic Model 3* (Fig. 6; Additional file 1: Table S8). The realistic model produced higher mean cancer-cell nuclear doses across all source localizations, and substantially broader cellular dose distributions, with some cancer cells receiving markedly lower doses than predicted by the uniform representation (Fig. 6a). The corresponding survival-fraction distributions at *A*_0_ = 0.1 Bq were similarly broader in the realistic geometry, particularly for ^103^Pd and ^161^Tb, ranging from near-complete killing to near-survival within the same cluster (Fig. 6b). A uniform intra-voxel activity assumption therefore obscures low-dose cancer-cell subpopulations that may remain insufficiently irradiated even when the mean voxel dose appears adequate.

**Fig. 6:**
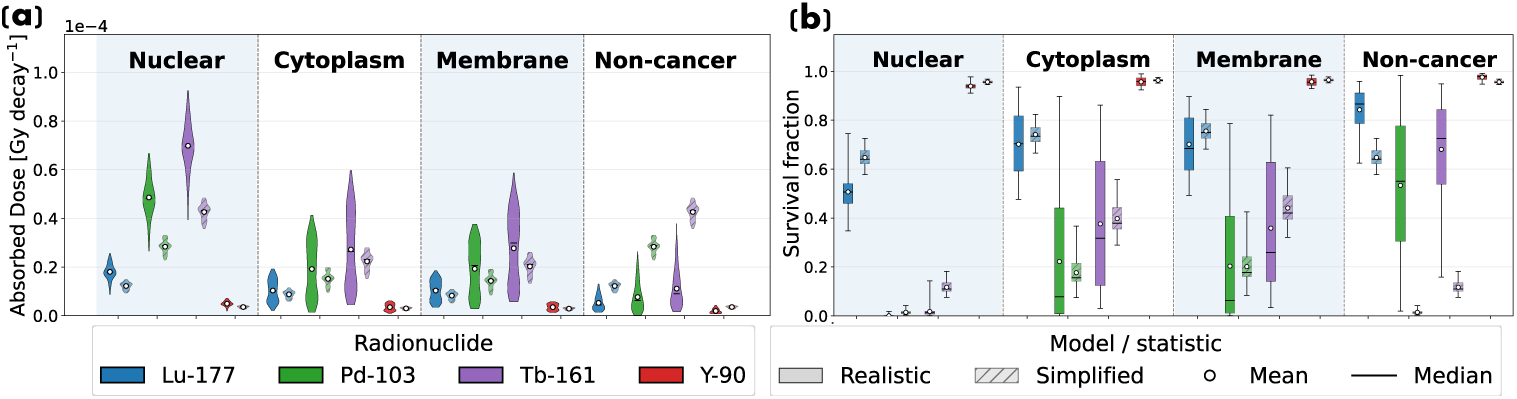
Comparison of cellular absorbed doses and predicted cancer-cell survival fractions in the uniformly labeled *Simplified Model 3* and the corresponding image-derived *Realistic Model 3* used to examine intra-voxel heterogeneity. (a) Mean absorbed dose to cancer-cell nuclei per cluster decay. (b) Mean cancer-cell survival fractions at an initial total cluster activity of *A*_0_ = 0.1 Bq. Column headers Nuclear, Cytoplasm, Membrane, and Non-cancer denote activity localized in the cancer-cell nucleus, cytoplasm, and membrane-associated compartment, and in non-cancer cells, respectively.

## 4 Discussion

We developed realistic 3D cellular tumor models from thick-section 3D CyCIF imaging to evaluate how native tumor microarchitecture affects cellular absorbed-dose distributions and predicted radiobiological response in TRT. Monte Carlo simulations of ^177^Lu, ^103^Pd, ^161^Tb and ^90^Y were performed with activity localized in the cancer-cell nucleus, cytoplasm, or membrane, and in non-cancer cells, across three model pairs. First, we compared a small 19-cell cluster represented using realistic and simplified geometries *(Realistic/Simplified Model 1)* to characterize the effect of cellular clustering and geometry in isolation (Fig. 2a and Fig. 2d). Next, we compared two larger tumor regions derived from the realistic data with their simplified counterparts, in which the same proportion of cells was labeled as cancer cells according to the cancer-cell fraction calculated from the corresponding realistic model (Fig. 2b, Fig. 2e and Fig. 2f). We also compared *Realistic Model 3* with the uniformly labeled *(Simplified Model 3)* to isolate the effect of intra-voxel heterogeneity. Finally, using *Realistic Model 3*, we examined cross-irradiation of cancer cells arising from activity localized in neighboring non-cancer cells.

The simplified large-cluster models were designed to represent how the corresponding realistic tissue regions might be approximated using conventional cellular-dosimetry frameworks such as MIRDcell [12], which is particularly useful for modeling idealized multicellular spheroids and tumor clusters, where cells are represented as regularly packed spheres with activity assigned to selected populations. Because MIRDcell does not reproduce the irregular cellular morphology, variable packing density, or spatial organization of cancer and non-cancer cells present in native tumor microarchitecture, the cancer-cell fraction measured from each image-derived region was used to randomly assign the equivalent fraction of spherical cells as radiolabeled cancer cells in the corresponding simplified model. This approach preserves the overall tissue volume and cancer-cell fraction while isolating the effect of detailed spatial organization on dosimetric and radiobiological outcomes.

Across both model types, absorbed dose per decay to cancer-cell nuclei ranked ^161^Tb *>* ^103^Pd *>*^177^Lu *>* ^90^Y consistent with cellular S-value calculations by Fourie et al. identifying ^161^Tb and ^103^Pd as high cellular-dose contributors relative to ^177^Lu [5]. In the 19-cell cluster comparison, *Simplified Model 1* overestimated mean nuclear absorbed dose relative to *Realistic Model 1* for every radionuclide and source localization examined, modestly for intranuclear activity (9.54% to 14.51%), but substantially for cytoplasmic and membrane-associated activity (33.23% to 44.88%), indicating that the simplified geometry most strongly overestimates dose when activity resides outside the nucleus. This agrees with earlier single-cell dosimetry studies: Tamborino et al. reported even larger overestimates for spherical MIRDcell-type geometries relative to microscopy-derived isolated cell models (65% for cytoplasmic and 42% for membrane source localization) [10], Tang et al. found comparable discrepancies that grew when source and target regions were spatially separated [21], and Arnaud et al. showed that realistic cell geometry and non-uniform source distributions alter energy deposition for ^125^I Auger-electron emitters [15]. The present study extends these findings from isolated cells to image-derived 3D cancer-cell clusters, where both cell morphology and local packing contribute to the absorbed-dose differences between simplified and realistic models.

We also found out that mean absorbed dose alone did not fully describe cellular irradiation in the larger model comparisons. In analysis including *Simplified Model 2* and *Realistic Model 2*, the simplified geometry reproduced the average nuclear doses reasonably well in several cases (Fig. 4a–b). A similar agreement was observed for mean cancer-cell nuclear dose in the more spatially heterogeneous *Realistic Model 3* (Fig. 4c). However, the corresponding cellular dose distributions differed substantially, with the realistic models consistently producing broader dose ranges that included both low-dose and high-dose subpopulations not captured by simplified models. Spatial organization also altered non-cancer-cell dosimetry; while simplified and realistic models produced broadly comparable mean non-cancer-cell doses in the intermixed region, the realistic model yielded consistently lower values in the more spatially heterogeneous *Realistic Model 3*, with reductions ranging from 34% to 47%, likely reflecting the spatial segregation of cancer and non-cancer cells, which increases the source– target distance for many non-cancer-cell nuclei. These effects were particularly evident for ^103^Pd and ^161^Tb, whose short-range electron emissions are more sensitive to local variations in cell spacing and source–target geometry. Although random labeling may provide a reasonable approximation of mean absorbed dose in an intermixed tumor region, it does not reproduce the structural and morphological heterogeneity of realistic tissue.

At a given initial cluster activity, ^103^Pd produced the greatest overall cancer-cell killing effect, followed by ^161^Tb, ^177^Lu and ^90^Y (Fig. 5). The therapeutic potential of ^103^Pd has also been highlighted by Hindié et al., who reported higher absorbed doses for ^103^Pd/^103^*^m^*Rh relative to ^161^Tb and ^177^Lu after energy normalization [4]. The broader absorbed-dose distributions in the realistic models translated directly into wider survival-fraction distributions, with subpopulations of cancer cells receiving insufficient dose and retaining relatively high survival probabilities in both the intermixed and more spatially heterogeneous tumor regions. Previous work also shows that spatially heterogeneous radionuclide distributions can strongly influence biological effect and tumor control probability, particularly when weakly irradiated cell populations remain within the tumor [22, 23]. These findings motivated the subsequent TCP analysis evaluating the combined effect of cell-level survival heterogeneity on the probability of controlling the entire modeled cancer-cell population.

The effect of these weakly irradiated cancer-cell subpopulations became more apparent in the TCP analysis of the more spatially heterogeneous *Realistic Model 3*. The realistic model required a higher initial cluster activity to reach *TCP*_90_ for every radionuclide and source-localization combination examined (Table 2). The differences were relatively small for nuclear localization but increased substantially when activity was localized in the cytoplasm or on the membrane. This is expected as nuclear localization provides a strong self-dose contribution, whereas extra-nuclear localization is more sensitive to variations in cell morphology, source-target distance, and the spatial arrangement of neighboring cells which was evident in the absorbed dose distributions. The greatest difference was observed for cytoplasmic localization of ^103^Pd, for which the realistic model required 5.93 times higher activity than the simplified model. Although ^103^Pd remained highly effective in terms of the absolute activity required to achieve tumor control, its predominantly short-range electron emissions made its efficacy more sensitive to spatial heterogeneity when the activity was localized outside the nucleus. Hindié et al., also showed that non-uniform targeting of ^103^Pd/^103^*^m^*Rh, produced substantial dose heterogeneity in unlabeled cells in their 19 cluster model [4]. These results extend previous studies showing that heterogeneous radionuclide distributions can strongly influence biological response and TCP, particularly when spatial non-uniformity produces weakly irradiated cancer-cell subpopulations.

Comparing treatment configurations for equal tumor control (*TCP*_90_) further underscores the influence of tumor microarchitecture on treatment evaluation. The simplified and image-derived models differed in the configuration associated with greatest non-cancer-cell preservation at *TCP*_90_ (Table 3): nuclear localization of ^161^Tb was most favorable in the simplified model, whereas nuclear localization of ^103^Pd was favored in the realistic model. These discrepancies suggest that treatment-related conclusions derived from idealized cellular geometries may not fully reflect the response predicted in heterogeneous tumor tissue, and in-silico studies relying solely on simplified cell models may therefore fail to capture the true radiobiological effects of a candidate radionuclide or targeting strategy. We therefore propose that incorporating histopathology-informed microarchitecture may provide a more representative basis for evaluating radionuclides and subcellular targeting strategies.

We also examined how targeting non-cancer cells, affects dose deposition and downstream radiobiological response. Deliberately targeting non-cancer cells within the tumor microenvironment has recently gained clinical traction as a treatment strategy. For example, Fibroblast activation protein (FAP) overexpressed on cancer-associated fibroblasts in the stroma of most epithelial malignancies while minimally expressed in normal tissue, has emerged as a pan-tumor target, with several FAP-targeted agents labeled with ^90^Y, ^177^Lu, and other radionuclides now in early-phase clinical trials [24]. 3D CyCIF data are highly valuable for investigating such strategies, as the technique separately stains distinct cell populations within the tumor microenvironment, cancer, immune, and other stromal cells, allowing source localization to be reassigned to any of these populations for simulation and radiobiological-effect estimation, as demonstrated in our non-cancer-cell cross-irradiation analysis.

Quantitative SPECT/PET imaging has become of paramount importance for the realization of accurate, patient-specific dosimetry in TRT [25, 26]. However, translating these macroscopic images into precise dose maps presents inherent challenges. As established by Ljungberg et al., the dispersion of absorbed dose measurements within a given volume of interest stems from four main sources: actual non-uniformities in tissue activity, external particle radiation cross-fire, partial-volume errors caused due to finite spatial resolution (spill-in and spill-out effects), and noise acquired during SPECT/PET data collection [26]. Considerable effort has been directed toward mitigating partial-volume errors through optimized reconstruction parameters and post-reconstruction correction methods. Although these studies address important imaging-related limitations, a more fundamental challenge remains: genuine microscopic activity heterogeneity within a voxel cannot be resolved with current clinical imaging systems, where voxel dimensions are constrained to several millimeters, and further reductions in voxel size are not presently feasible [9, 25]. We investigated how realistic non-uniform activity distributions at the sub-voxel scale influence absorbed doses to target nuclei and the resulting radiobiological response by comparing uniformly labeled *Simplified Model 3* with the corresponding image-derived realistic tissue model. The realistic model consistently produced higher mean absorbed doses to cancer-cell nuclei across all radionuclides and sub-cellular source localizations (Fig. 6a). This outcome is physically intuitive: while an imaging voxel may appear to contain uniformly distributed activity, that activity may in reality be concentrated within a small subset of cells, giving rise to higher local absorbed doses. For non-cancer-cell localization, by contrast, the realistic model produced lower mean cross-dose to cancer-cell nuclei than the simplified model. In *Simplified Model 3*, the nominal non-cancer-cell-localization condition reduces to activity distributed among the cancer cells themselves, considering a uniform activity distribution inside an imaging voxel, approximating a self-dose scenario. In *Realistic Model 3*, however, non-cancer cells occupy a spatially distinct compartment, so cancer-cell nuclei receive dose exclusively through genuine cross-irradiation from neighboring non-cancer cells. The observed heterogeneity in dose distribution has direct radiobiological consequences, as reflected in the survival fraction distribution (Fig. 6b). It can be seen that a subset of cells may receive sublethal doses, potentially compromising treatment efficacy. This is consistent with our earlier findings, which demonstrated that intra-cluster activity heterogeneity can substantially alter predicted cell survival fractions and, by extension, influence treatment decisions. Collectively, these results underscore that sub-voxel activity heterogeneity carries genuine radiobiological significance that extends well beyond the reach of current imaging resolution, and future patient-specific dosimetry frameworks should incorporate strategies to account for these effects if they are to provide dosimetric estimates that are truly predictive of treatment outcome.

Several limitations should be considered when interpreting the present findings. First, fixed radiosensitivity parameters were used to calculate survival fractions for cancer and non-cancer cells. In reality, radiosensitivity may vary across cancer phenotypes, normal-cell types, radionuclides, dose rates, and subcellular source localizations. The non-cancer-cell population, which may include immune, stromal, endothelial, and other tissue-associated cell types, was represented using a common reference parameter set despite its cellular diversity. Therefore, the predicted non-cancer-cell survival fractions should be interpreted as comparative indicators rather than direct estimates of normal-tissue toxicity. An important advantage of multiplexed 3D CyCIF imaging is that additional cell-type-specific markers can be incorporated to distinguish these different cellular populations. Future studies could therefore evaluate absorbed dose and predicted survival separately for individual non-cancer-cell types, including immune and other stromal cells.

Activity was assumed to be uniformly distributed within the assigned cellular compartment. In practice, radiopharmaceutical uptake is governed by target expression, ligand binding, internalization, intracellular trafficking, and retention kinetics. For example, in PSMA-targeted radiopharmaceutical therapy, the radiolabeled ligand binds to prostate-specific membrane antigen (PSMA) expressed on the cancer-cell surface and may subsequently undergo partial internalization. These processes could not be resolved from the present imaging dataset. Future studies could incorporate spatially resolved molecular information to better characterize target-expression patterns and inform more realistic activity distributions. In particular, the integration of 3D CyCIF imaging with spatial transcriptomic or proteomic data may enable the development of cell-specific radiopharmaceutical binding models and support more detailed evaluation of downstream absorbed-dose deposition and biological response.

The simplified and realistic large cluster models did not contain identical numbers of cancer cells, despite representing comparable overall tissue volumes. We envision this difference as an inherent consequence of realistic tissue morphology, arising from variations in cell size, irregular cellular shape, packing density, and heterogeneous tissue composition within the image-derived models. However, it should be acknowledged that TCP depends on both the survival probabilities of individual cancer cells and the total number of cancer cells within the modeled region. Consequently, differences in cancer-cell count may contribute to variations in the predicted TCP alongside the effects of cellular dose heterogeneity.

3D CyCIF relies on iterative cycles of antibody staining, imaging, and fluorophore inactivation, and repeated processing of this kind has been reported to progressively degrade tissue integrity. Yapp et al. report that although most specimens tolerated this procedure well, a subset of samples disintegrated during antigen retrieval, and separately documented a ”corrugation” artifact. Because the realistic cellular models in this study were constructed directly from segmented CyCIF volumes, any such processing-related distortion of nuclear size, cytoplasmic thickness, or intercellular spacing would propagate directly into the simulated dose distributions. Future work applying this pipeline to additional specimens, ideally with paired assessment of tissue integrity, would help establish how sensitive the reported doses are to these preanalytical factors.

We also note that the workflow presented here, requiring thick-section multiplexed volumetric imaging together with per-cell Monte Carlo radiation transport simulation, is computationally and logistically intensive and is not presently suited to routine clinical dosimetry. Generating a single realistic tumor phantom requires specialized tissue acquisition, multi-day multiplexed imaging protocols, and substantial downstream image segmentation and simulation runtime. We therefore envision the present framework primarily as a research and validation tool, that can be helpful in benchmarking and calibrating radionuclide development for upcoming new radionuclides. As multiplexed imaging technologies and computational pipelines continue to mature, we anticipate that histopathology-informed dosimetry will become an increasingly accessible component of preclinical radionuclide evaluation, helping to prioritize the most promising candidates and subcellular targeting strategies before they advance to costly in vivo and clinical testing.

## 5 Conclusion

We demonstrated that native 3D tumor microarchitecture can substantially influence cellular-scale dosimetry and predicted radiobiological response in TRT. Histopathology-informed models derived from 3D CyCIF imaging revealed broader cancer and non-cancer-cell absorbed-dose distributions than simplified spherical models, including low-dose cancer-cell subpopulations that were not captured by idealized geometries. These differences propagated into wider survival-fraction distributions and altered the activity required to achieve tumor control, particularly for cytoplasmic and membrane source localization. It was also found out that treatment related conclusions may be sensitive to the underlying tissue geometry. The intra-voxel analysis further showed that voxel-averaged representations may obscure biologically relevant microscopic heterogeneity. Overall, these findings support the use of histopathology-informed 3D models as a complementary framework for improving the interpretation of cellular dosimetry and refining radionuclide and subcellular targeting strategies.

## Supplementary information

Supplementary material accompanies this article. **Additional file 1: Table S1a**: geometrical and cellular properties of *Simplified Model 1* and *Realistic Model 1* ; **Table S1b**: the same for *Simplified* and *Realistic Models 2* and *3*. **Table S2**: mean percentage Monte Carlo dose uncertainty per cancer-cell nucleus, by radionuclide and subcellular source localization (10^6^ simulated decays per condition). **Table S3**: mean cancer-cell nuclear absorbed dose per cluster decay for *Simplified Model 1* and *Realistic Model 1*, with statistical comparison (two-sided one-sample *t*-test). **Table S4a** and **Table S4b**: mean absorbed dose to cancer-cell and non-cancer-cell nuclei per cluster decay, respectively, for *Simplified Model 2* and *Realistic Model 2* ; **Fig. S2** and **Fig. S3**: the corresponding mean cancer-cell and non-cancer-cell nuclear doses for *Realistic Model 2* and *Simplified Model 2*. **Table S5a** and **Table S5b**: mean absorbed dose to cancer-cell and non-cancer-cell nuclei per cluster decay for *Simplified Model 2* and *Realistic Model 3* ; **Fig. S4** and **Fig. S5**: the corresponding mean cancer-cell and non-cancer-cell nuclear doses for *Realistic Model 3* and *Simplified Model 2*. **Table S6a** and **Table S6b**: mean cancer-cell and non-cancer-cell survival fractions for *Simplified Model 2* and *Realistic Model 2* at 0.08 Bq. **Table S7a** and **Table S7b**: mean cancer-cell and non-cancer-cell survival fractions for *Simplified Model 2* and *Realistic Model 3* at 0.1 Bq. **Table S8**: mean cancer-cell nuclear absorbed dose per total cluster decay for *Simplified Model 3* and *Realistic Model 3*, characterizing intra-voxel dose heterogeneity; **Fig. S6**: the corresponding mean cellular nuclear dose for *Realistic Model 3* and *Simplified Model 3*. **Table S9**: physical decay characteristics of the radionuclides used in this study.

## Abbreviations

3D: Three-dimensional
CyCIF: Cyclic immunofluorescence
FAP: Fibroblast activation protein
GATE: Geant4 Application for Tomographic Emission
MIRD: Medical Internal Radiation Dose
PET: Positron emission tomography
PSMA: Prostate-specific membrane antigen
SF: Survival fraction
SPECT: Single-photon emission computed tomography
TCP: Tumor control probability
TIAC: Time-integrated activity coefficient
TRT: Targeted radionuclide therapy
VGP: Vertical growth phase

## Declarations

### Ethics approval and consent to participate

Not applicable. This study did not involve any new experiments on human participants, human tissue, or animals. All analyses were performed on a previously published, publicly available, de-identified 3D CyCIF imaging dataset [27].

### Consent for publication

Not applicable.

### Availability of data and materials

The dataset supporting the conclusions of this article is available in the Zenodo repository, https://doi.org/10.5281/zenodo.10055593 [27]. The analysis and simulation code generated during the current study is available in the GitHub repository, https://github.com/TreshanAyesh/Cycif-dose.

### Competing interests

RW is a founder and the Clinical Director of Cyclotek (Aust) Pty Ltd. The remaining authors declare no competing interests.

### Funding

The PhD candidature of TP is partly funded by Cyclotek (Aust) Pty Ltd.

### Authors’ contributions

TP contributed to methodology, experimentation, analysis, and writing of the manuscript. RW contributed to concept formulation, analysis, and editing. RT, SK, PHM and JT contributed to concept formulation and editing. All authors read and approved the final manuscript.

## Acknowledgements

Not applicable.

## Notes

### Competing Interest Statement

Dr. Robert Ware is a founder and the Clinical Director of Cyclotek (Aust) Pty Ltd, which partly funds the PhD scholarship of the first author, Treshan Peirispulle. The remaining authors declare no competing interests.

