## Additional file 1 for "Impact of Realistic 3D Tumor Microarchitecture on Cellular Dosimetry and Radiobiological Response in Targeted Radionuclide Therapy"

#### Supplementary Materials

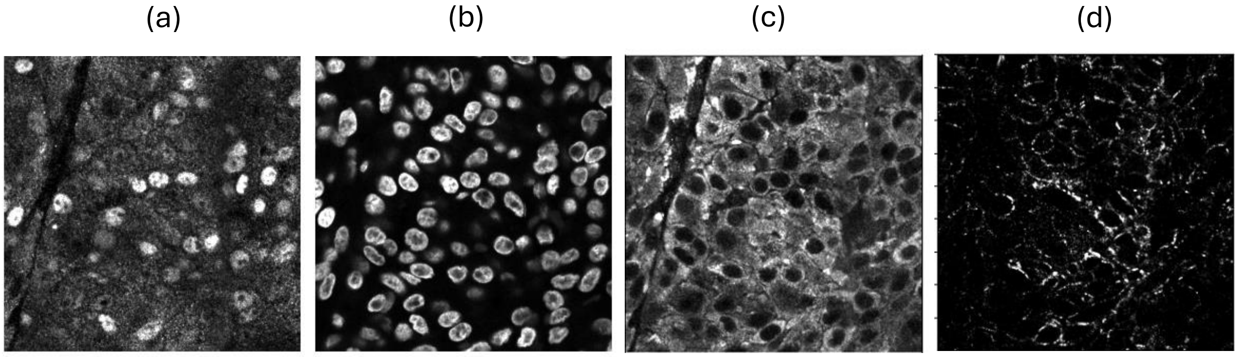


*Fig. S1.* *Representative 3D CyCIF channels used for cellular segmentation. (a) SOX10 channel, identifying cancer-cell nuclei. (b) Hoechst channel, marking all nuclei (DNA). (c) MART1 channel, delineating cancer-cell cytoplasm. (d) β - catenin channel, shown for reference; membrane-associated regions were defined computationally as the boundary voxels of the segmented cytoplasm mask (see Methods).*

### Supplementary Table S1

##### Table S1a. Geometrical and cellular properties of Simplified Model 1 and Realistic Model 1.

| Parameter | Simplified Model 1 | Realistic Model 1 |
|---|---|---|
| **Geometry type** | Idealized concentric spheres | Image-derived (3D CyCIF) |
| **Total cells** | 19 | 19 per cluster (n = 20 clusters) |
| **Cancer cells** | 19 | 19 |
| **Non-cancer cells** | 0 | 19.3 ± 7.0 |
| **Mean cell diameter (μm)** | 10 | 10.38 ± 2.79 |
| **Mean nuclear diameter (μm)** | 8 | 7.95 ± 1.53 |
| **Spatial arrangement** | Regular: 1 central + 6 first neighbor + 12 second-neighbor cells | Native, irregular packing derived from tissue |

##### Table S1b. Geometrical and cellular properties of Simplified and Realistic Models 2 and 3.

| Parameter | Simplified Model 2 | Realistic Model 2 | Simplified Model 3 | Realistic Model 3 |
|---|---|---|---|---|
| **Geometry type** | Idealized sphere | Image-derived | Idealized sphere | Image-derived |
| **Overall dimensions** | r = 61 μm, h = 30 μm | r = 61 μm, h = 30 μm | r = 61 μm, h = 30 μm | r = 61 μm, h = 30 μm |
| **Total cells** | 335 | 250 | 335 | 364 |
| **Cancer cells** | † | 118 | 335 | 183 |
| **Non-cancer cells** | † | 132 | 0 | 181 |
| **Cancer-cell fraction (%)** | † | 47.2 | 100 | 50.2 |
| **Spatial arrangement** | Regular, randomly labeled | Native, intermixed | Regular, uniformly labeled (all cancer) | Native, spatially concentrated subregion |

*Note: Realistic Model 1 statistics reflect the mean across 20 independently sampled clusters (see Methods §2.4).* † *Number of cancer cells and non-cancer cells are calculated according to the cancer-cell fraction of the corresponding realistic model in comparison.*

### Supplementary Table S2

**Table S2. Mean percentage Monte Carlo dose uncertainty per cancer-cell nucleus, by radionuclide and subcellular source localization ($10^6$ simulated decays per condition).**

| Radionuclide | Source localization | Mean MC dose uncertainty (%) |
|---|---|---|
| Lu-177 | Nucleus | 1.03 |
| Lu-177 | Cytoplasm | 1.06 |
| Lu-177 | Membrane | 1.02 |
| Lu-177 | Non-cancer cell | 0.85 |
| Pd-103 | Nucleus | 1.85 |
| Pd-103 | Cytoplasm | 1.90 |
| Pd-103 | Membrane | 1.52 |
| Pd-103 | Non-cancer cell | 1.18 |
| Tb-161 | Nucleus | 0.89 |
| Tb-161 | Cytoplasm | 0.88 |
| Tb-161 | Membrane | 0.83 |
| Tb-161 | Non-cancer cell | 0.71 |
| Y-90 | Nucleus | 1.46 |
| Y-90 | Cytoplasm | 1.53 |
| Y-90 | Membrane | 1.48 |
| Y-90 | Non-cancer cell | 1.23 |

*Note: "Non-cancer cell" localization corresponds to the cross-irradiation scenario in which activity was localized within the non-cancer-cell population (see §2.4, §3.5).*

### Supplementary Table S3

**Table S3. Mean cancer-cell nuclear absorbed dose per cluster decay for Simplified Model 1 and Realistic Model 1, with statistical comparison (two-sided one-sample t-test), by radionuclide and subcellular source localization. Corresponds to Fig. 3.**

| Radionuclide | Source localization | Simplified Model 1 dose ± SD ($\times 10^{-4}$ Gy; n = 19 cells) | Realistic Model 1 mean ± SD ($\times 10^{-4}$ Gy; n = 20 clusters) | Relative difference (%) | t-statistic | p-value |
|---|---|---|---|---|---|---|
| Lu-177 | Nucleus | 1.30 ± 0.07 | *1.11 ± 0.12* | -14.51 | -4.79 | $9.90 \times 10^{-4}$ |
| Lu-177 | Cytoplasm | 0.68 ± 0.08 | *0.40 ± 0.06* | -40.67 | -15.73 | $7.48 \times 10^{-8}$ |
| Lu-177 | Membrane | 0.59 ± 0.08 | *0.39 ± 0.07* | -33.23 | -9.02 | $8.40 \times 10^{-6}$ |
| Pd-103 | Nucleus | 4.03 ± 0.22 | *3.64 ± 0.43* | -9.54 | -2.84 | 0.0194 |
| Pd-103 | Cytoplasm | 1.65 ± 0.23 | *0.92 ± 0.14* | -44.09 | -16.04 | $6.31 \times 10^{-8}$ |
| Pd-103 | Membrane | 1.47 ± 0.23 | *0.90 ± 0.17* | -38.95 | -10.45 | $2.47 \times 10^{-6}$ |
| Tb-161 | Nucleus | 5.94 ± 0.22 | *5.23 ± 0.62* | -11.91 | -3.62 | 0.0056 |
| Tb-161 | Cytoplasm | 2.28 ± 0.24 | *1.26 ± 0.20* | -44.88 | -16.25 | $5.63 \times 10^{-8}$ |
| Tb-161 | Membrane | 1.88 ± 0.25 | *1.21 ± 0.24* | -35.52 | -8.64 | $1.19 \times 10^{-5}$ |
| Y-90 | Nucleus | 0.31 ± 0.02 | *0.26* ± 0.03 | -15.86 | -5.37 | $4.52 \times 10^{-4}$ |
| Y-90 | Cytoplasm | 0.20 ± 0.02 | *0.13* ± 0.02 | -37.62 | -14.55 | $1.47 \times 10^{-7}$ |
| Y-90 | Membrane | 0.18 ± 0.02 | *0.12* ± 0.03 | -33.05 | -9.73 | $4.47 \times 10^{-6}$ |

*Note: Simplified Model 1 mean and SD reflect the dose distribution across the 19 cancer cells within the single simplified cluster. Realistic Model 1 mean is the average of the per-cluster mean doses, and SD is the average of the within-cluster standard deviations, across the sampled clusters. For each radionuclide and source-localization combination, a two-sided one-sample t-test compared the fixed Simplified Model 1 dose value against the distribution of Realistic Model 1 cluster dose values; all comparisons were statistically significant ($p < 0.05$). Relative difference is calculated as (Realistic − Simplified) / Simplified × 100; negative values indicate that the Simplified Model overestimated dose relative to the Realistic Model.*

### Supplementary Table S4

**Table S4a. Mean absorbed dose to cancer-cell nuclei per cluster decay for Simplified Model 2 and Realistic Model 2.**

| Radionuclide | Compartment | Mean dose (Simp.) [Gy decay$^{-1}$] | Mean dose (Real.) [Gy decay$^{-1}$] | SD (Simp.) [Gy decay$^{-1}$] | SD (Real.) [Gy decay$^{-1}$] | Diff. (%) |
|---|---|---|---|---|---|---|
| Lu-177 | Nucleus | $1.80 \times 10^{-5}$ | $2.06 \times 10^{-5}$ | $1.59 \times 10^{-6}$ | $3.09 \times 10^{-6}$ | +14.1 |
| Pd-103 | Nucleus | $4.80 \times 10^{-5}$ | $6.07 \times 10^{-5}$ | $3.22 \times 10^{-6}$ | $7.40 \times 10^{-6}$ | +26.5 |
| Tb-161 | Nucleus | $7.50 \times 10^{-5}$ | $8.97 \times 10^{-5}$ | $3.78 \times 10^{-6}$ | $1.08 \times 10^{-5}$ | +19.60 |
| Y-90 | Nucleus | $5.00 \times 10^{-6}$ | $5.53 \times 10^{-6}$ | $5.45 \times 10^{-7}$ | $9.50 \times 10^{-7}$ | +10.64 |
| Lu-177 | Cytoplasm | $1.10 \times 10^{-5}$ | $9.20 \times 10^{-6}$ | $1.62 \times 10^{-6}$ | $2.33 \times 10^{-6}$ | -16.35 |
| Pd-103 | Cytoplasm | $2.00 \times 10^{-5}$ | $1.65 \times 10^{-5}$ | $3.33 \times 10^{-6}$ | $5.49 \times 10^{-6}$ | -17.24 |
| Tb-161 | Cytoplasm | $3.00 \times 10^{-5}$ | $2.29 \times 10^{-5}$ | $4.02 \times 10^{-6}$ | $7.22 \times 10^{-6}$ | -23.62 |
| Y-90 | Cytoplasm | $4.00 \times 10^{-6}$ | $3.50 \times 10^{-6}$ | $5.57 \times 10^{-7}$ | $7.38 \times 10^{-7}$ | -12.43 |
| Lu-177 | Membrane | $1.00 \times 10^{-5}$ | $9.09 \times 10^{-6}$ | $1.67 \times 10^{-6}$ | $2.03 \times 10^{-6}$ | -9.09 |
| Pd-103 | Membrane | $1.80 \times 10^{-5}$ | $1.60 \times 10^{-5}$ | $3.40 \times 10^{-6}$ | $4.46 \times 10^{-6}$ | -10.87 |
| Tb-161 | Membrane | $2.60 \times 10^{-5}$ | $2.28 \times 10^{-5}$ | $4.10 \times 10^{-6}$ | $5.94 \times 10^{-6}$ | -12.44 |
| Y-90 | Membrane | $3.00 \times 10^{-6}$ | $2.77 \times 10^{-6}$ | $5.61 \times 10^{-7}$ | $6.67 \times 10^{-7}$ | -7.64 |
| Lu-177 | Non-cancer | $7.00 \times 10^{-6}$ | $6.32 \times 10^{-6}$ | $1.68 \times 10^{-6}$ | $1.60 \times 10^{-6}$ | -9.73 |
| Pd-103 | Non-cancer | $1.10 \times 10^{-5}$ | $9.59 \times 10^{-6}$ | $3.54 \times 10^{-6}$ | $3.64 \times 10^{-6}$ | -12.86 |
| Tb-161 | Non-cancer | $1.40 \times 10^{-5}$ | $1.28 \times 10^{-5}$ | $4.05 \times 10^{-6}$ | $4.38 \times 10^{-6}$ | -8.83 |
| Y-90 | Non-cancer | $2.00 \times 10^{-6}$ | $1.83 \times 10^{-6}$ | $5.71 \times 10^{-7}$ | $5.34 \times 10^{-7}$ | -8.64 |

**Table S4b. Mean absorbed dose to non-cancer-cell nuclei per cluster decay for Simplified Model 2 and Realistic Model 2.**

| Radionuclide | Compartment | Mean dose (Simp.) [Gy decay$^{-1}$] | Mean dose (Real.) [Gy decay$^{-1}$] | SD (Simp.) [Gy decay$^{-1}$] | SD (Real.) [Gy decay$^{-1}$] | Diff. (%) |
|---|---|---|---|---|---|---|
| Lu-177 | Nucleus | $7.00 \times 10^{-6}$ | $6.16 \times 10^{-6}$ | $1.66 \times 10^{-6}$ | $2.34 \times 10^{-6}$ | -11.9% |
| Pd-103 | Nucleus | $1.10 \times 10^{-5}$ | $9.35 \times 10^{-6}$ | $3.37 \times 10^{-6}$ | $5.24 \times 10^{-6}$ | -15.0% |
| Tb-161 | Nucleus | $1.40 \times 10^{-5}$ | $1.25 \times 10^{-5}$ | $3.92 \times 10^{-6}$ | $6.30 \times 10^{-6}$ | -10.7% |
| Y-90 | Nucleus | $3.00 \times 10^{-6}$ | $2.66 \times 10^{-6}$ | $5.64 \times 10^{-7}$ | $7.77 \times 10^{-7}$ | -11.5% |
| Lu-177 | Cytoplasm | $7.00 \times 10^{-6}$ | $6.67 \times 10^{-6}$ | $1.71 \times 10^{-6}$ | $2.87 \times 10^{-6}$ | -4.7% |
| Pd-103 | Cytoplasm | $1.10 \times 10^{-5}$ | $1.12 \times 10^{-5}$ | $3.49 \times 10^{-6}$ | $7.64 \times 10^{-6}$ | +1.8% |
| Tb-161 | Cytoplasm | $1.50 \times 10^{-5}$ | $1.60 \times 10^{-5}$ | $4.19 \times 10^{-6}$ | $1.03 \times 10^{-5}$ | +6.4% |
| Y-90 | Cytoplasm | $3.00 \times 10^{-6}$ | $2.77 \times 10^{-6}$ | $5.73 \times 10^{-7}$ | $8.55 \times 10^{-7}$ | -7.6% |
| Lu-177 | Membrane | $7.00 \times 10^{-6}$ | $7.06 \times 10^{-6}$ | $1.71 \times 10^{-6}$ | $3.03 \times 10^{-6}$ | +0.8% |
| Pd-103 | Membrane | $1.10 \times 10^{-5}$ | $1.24 \times 10^{-5}$ | $3.56 \times 10^{-6}$ | $7.91 \times 10^{-6}$ | +12.8% |
| Tb-161 | Membrane | $1.60 \times 10^{-5}$ | $1.87 \times 10^{-5}$ | $4.33 \times 10^{-6}$ | $1.07 \times 10^{-5}$ | +16.8% |
| Y-90 | Membrane | $3.00 \times 10^{-6}$ | $2.90 \times 10^{-6}$ | $5.77 \times 10^{-7}$ | $9.02 \times 10^{-7}$ | -3.4% |
| Lu-177 | Non-cancer | $1.70 \times 10^{-5}$ | $1.85 \times 10^{-5}$ | $1.41 \times 10^{-6}$ | $2.54 \times 10^{-6}$ | +8.8% |
| Pd-103 | Non-cancer | $4.40 \times 10^{-5}$ | $5.26 \times 10^{-5}$ | $2.85 \times 10^{-6}$ | $5.96 \times 10^{-6}$ | +19.6% |
| Tb-161 | Non-cancer | $6.80 \times 10^{-5}$ | $7.65 \times 10^{-5}$ | $3.36 \times 10^{-6}$ | $8.45 \times 10^{-6}$ | +12.4% |
| Y-90 | Non-cancer | $5.00 \times 10^{-6}$ | $5.39 \times 10^{-6}$ | $4.74 \times 10^{-7}$ | $7.68 \times 10^{-7}$ | +7.8% |

*Simp. = Simplified Model 2; Real. = Realistic Model 2. SD = standard deviation across individual cells. Compartment indicates the cellular source localization. . Relative difference is calculated as (Realistic − Simplified) / Simplified × 100; negative values indicate that the Simplified Model overestimated dose relative to the Realistic Model.*

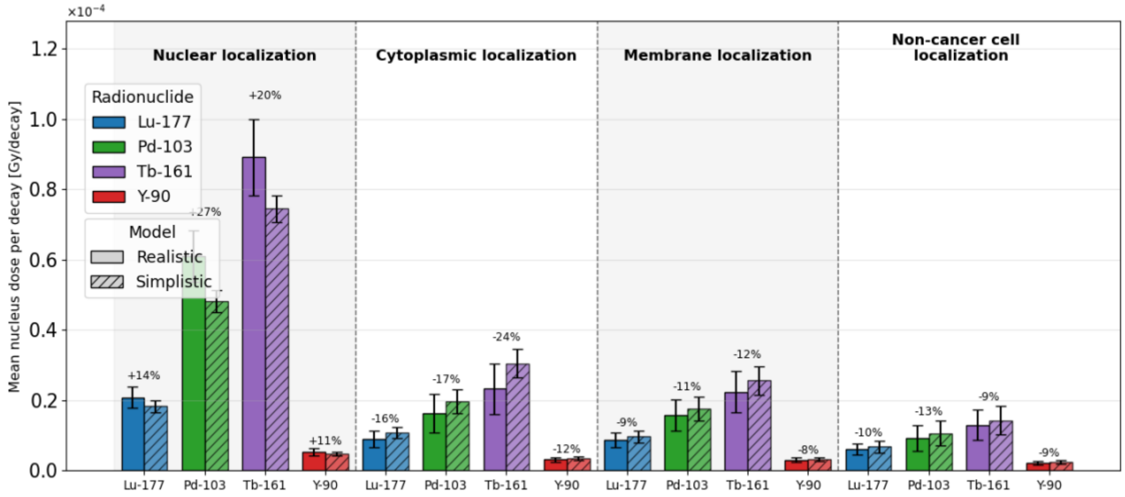


***Fig. S2.*** *Mean cancer-cell nuclear dose per cluster decay comparing Simplified Model 2 and Realistic Model 2. Solid bars = Realistic Model 2; hatched bars = Simplified Model 2. Error bars indicate the standard deviation across individual cells. Percentage labels denote the relative difference in mean dose for the realistic model compared with the simplified model.*

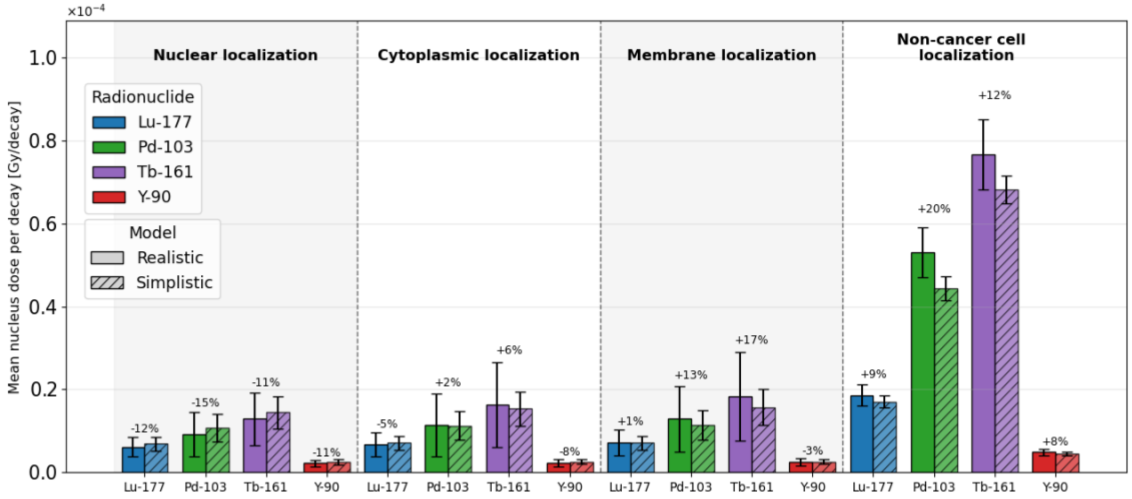


***Fig. S3.*** *Mean non-cancer-cell nuclear dose per cluster decay comparing Simplified Model 2 and Realistic Model 2. Solid bars = Realistic Model 2; hatched bars = Simplified Model 2. Error bars indicate the standard deviation across individual cells. Percentage labels denote the relative difference in mean dose for the realistic model compared with the simplified model.*

### Supplementary Table S5

**Table S5a. Mean absorbed dose to cancer-cell nuclei per cluster decay for Simplified Model 2 and Realistic Model 3.**

| Radionuclide | Compartment | Mean dose (Simp.) [Gy decay$^{-1}$] | Mean dose (Real.) [Gy decay$^{-1}$] | SD (Simp.) [Gy decay$^{-1}$] | SD (Real.) [Gy decay$^{-1}$] | Diff. (%) |
|---|---|---|---|---|---|---|
| Lu-177 | Nucleus | $1.80 \times 10^{-5}$ | $1.80 \times 10^{-5}$ | $1.53 \times 10^{-6}$ | $3.23 \times 10^{-6}$ | +2.2% |
| Pd-103 | Nucleus | $4.60 \times 10^{-5}$ | $4.86 \times 10^{-5}$ | $3.17 \times 10^{-6}$ | $7.67 \times 10^{-6}$ | +5.4% |
| Tb-161 | Nucleus | $7.10 \times 10^{-5}$ | $6.98 \times 10^{-5}$ | $3.70 \times 10^{-6}$ | $9.71 \times 10^{-6}$ | -1.8% |
| Y-90 | Nucleus | $5.00 \times 10^{-6}$ | $4.95 \times 10^{-6}$ | $5.25 \times 10^{-7}$ | $1.04 \times 10^{-6}$ | +6.7% |
| Lu-177 | Cytoplasm | $1.10 \times 10^{-5}$ | $1.03 \times 10^{-5}$ | $1.58 \times 10^{-6}$ | $4.50 \times 10^{-6}$ | -2.3% |
| Pd-103 | Cytoplasm | $1.90 \times 10^{-5}$ | $1.92 \times 10^{-5}$ | $3.31 \times 10^{-6}$ | $1.11 \times 10^{-5}$ | -0.6% |
| Tb-161 | Cytoplasm | $3.00 \times 10^{-5}$ | $2.72 \times 10^{-5}$ | $3.89 \times 10^{-6}$ | $1.53 \times 10^{-5}$ | -8.2% |
| Y-90 | Cytoplasm | $3.00 \times 10^{-6}$ | $3.49 \times 10^{-6}$ | $5.24 \times 10^{-7}$ | $1.38 \times 10^{-6}$ | +0.9% |
| Lu-177 | Membrane | $1.00 \times 10^{-5}$ | $1.03 \times 10^{-5}$ | $1.59 \times 10^{-6}$ | $4.13 \times 10^{-6}$ | +8.3% |
| Pd-103 | Membrane | $1.70 \times 10^{-5}$ | $1.93 \times 10^{-5}$ | $3.36 \times 10^{-6}$ | $1.02 \times 10^{-5}$ | +10.9% |
| Tb-161 | Membrane | $2.50 \times 10^{-5}$ | $2.77 \times 10^{-5}$ | $4.02 \times 10^{-6}$ | $1.42 \times 10^{-5}$ | +10.2% |
| Y-90 | Membrane | $3.00 \times 10^{-6}$ | $3.49 \times 10^{-6}$ | $5.34 \times 10^{-7}$ | $1.26 \times 10^{-6}$ | +7.4% |
| Lu-177 | Non-cancer | $7.00 \times 10^{-6}$ | $5.26 \times 10^{-6}$ | $1.74 \times 10^{-6}$ | $2.83 \times 10^{-6}$ | -22.7% |
| Pd-103 | Non-cancer | $1.10 \times 10^{-5}$ | $7.67 \times 10^{-6}$ | $3.67 \times 10^{-6}$ | $5.85 \times 10^{-6}$ | -27.7% |
| Tb-161 | Non-cancer | $1.40 \times 10^{-5}$ | $1.11 \times 10^{-5}$ | $4.20 \times 10^{-6}$ | $7.54 \times 10^{-6}$ | -21.6% |
| Y-90 | Non-cancer | $2.00 \times 10^{-6}$ | $1.95 \times 10^{-6}$ | $5.91 \times 10^{-7}$ | $9.34 \times 10^{-7}$ | -21.1% |

**Table S5b. Mean absorbed dose to non-cancer-cell nuclei per cluster decay for Simplified Model 2 and Realistic Model 3.**

| Radionuclide | Compartment | Mean dose (Simp.) [Gy decay$^{-1}$] | Mean dose (Real.) [Gy decay$^{-1}$] | SD (Simp.) [Gy decay$^{-1}$] | SD (Real.) [Gy decay$^{-1}$] | Diff. (%) |
|---|---|---|---|---|---|---|
| Lu-177 | Nucleus | $7.00 \times 10^{-6}$ | $4.57 \times 10^{-6}$ | $1.59 \times 10^{-6}$ | $2.98 \times 10^{-6}$ | -33.4% |
| Pd-103 | Nucleus | $1.10 \times 10^{-5}$ | $6.08 \times 10^{-6}$ | $3.21 \times 10^{-6}$ | $6.15 \times 10^{-6}$ | -42.8% |
| Tb-161 | Nucleus | $1.40 \times 10^{-5}$ | $9.25 \times 10^{-6}$ | $3.75 \times 10^{-6}$ | $7.86 \times 10^{-6}$ | -35.4% |
| Y-90 | Nucleus | $2.00 \times 10^{-6}$ | $1.72 \times 10^{-6}$ | $5.43 \times 10^{-7}$ | $9.97 \times 10^{-7}$ | -31.0% |
| Lu-177 | Cytoplasm | $7.00 \times 10^{-6}$ | $4.64 \times 10^{-6}$ | $1.65 \times 10^{-6}$ | $2.88 \times 10^{-6}$ | -33.8% |
| Pd-103 | Cytoplasm | $1.10 \times 10^{-5}$ | $5.96 \times 10^{-6}$ | $3.33 \times 10^{-6}$ | $6.33 \times 10^{-6}$ | -46.1% |
| Tb-161 | Cytoplasm | $1.50 \times 10^{-5}$ | $9.14 \times 10^{-6}$ | $4.03 \times 10^{-6}$ | $7.89 \times 10^{-6}$ | -39.7% |
| Y-90 | Cytoplasm | $3.00 \times 10^{-6}$ | $1.73 \times 10^{-6}$ | $5.62 \times 10^{-7}$ | $9.34 \times 10^{-7}$ | -31.8% |
| Lu-177 | Membrane | $7.00 \times 10^{-6}$ | $4.98 \times 10^{-6}$ | $1.64 \times 10^{-6}$ | $2.84 \times 10^{-6}$ | -29.8% |
| Pd-103 | Membrane | $1.10 \times 10^{-5}$ | $6.95 \times 10^{-6}$ | $3.37 \times 10^{-6}$ | $6.75 \times 10^{-6}$ | -38.1% |
| Tb-161 | Membrane | $1.60 \times 10^{-5}$ | $1.05 \times 10^{-5}$ | $4.13 \times 10^{-6}$ | $8.60 \times 10^{-6}$ | -32.6% |
| Y-90 | Membrane | $3.00 \times 10^{-6}$ | $1.85 \times 10^{-6}$ | $5.61 \times 10^{-7}$ | $9.15 \times 10^{-7}$ | -27.4% |
| Lu-177 | Non-cancer | $1.80 \times 10^{-5}$ | $2.17 \times 10^{-5}$ | $1.46 \times 10^{-6}$ | $3.25 \times 10^{-6}$ | +22.4% |
| Pd-103 | Non-cancer | $4.60 \times 10^{-5}$ | $6.20 \times 10^{-5}$ | $2.91 \times 10^{-6}$ | $7.86 \times 10^{-6}$ | +33.6% |
| Tb-161 | Non-cancer | $7.10 \times 10^{-5}$ | $8.85 \times 10^{-5}$ | $3.43 \times 10^{-6}$ | $1.04 \times 10^{-5}$ | +23.9% |
| Y-90 | Non-cancer | $5.00 \times 10^{-6}$ | $5.80 \times 10^{-6}$ | $4.88 \times 10^{-7}$ | $1.04 \times 10^{-6}$ | +23.9% |

*Simp. = Simplified Model 2; Real. = Realistic Model 3. SD = standard deviation across individual cells. Compartment indicates the source localization of the cancer-cell-targeted radiopharmaceutical. Diff. = percentage difference of the realistic relative to the simplified mean dose.*

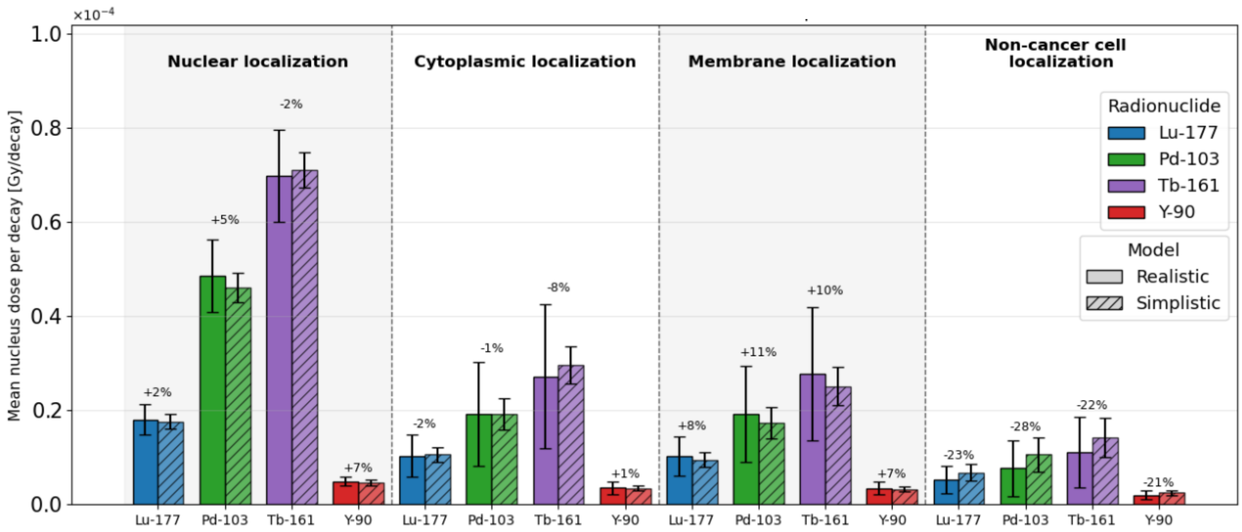


***Fig. S4.*** *Mean cancer-cell nuclear dose per cluster decay comparing Simplified Model 2 and Realistic Model 3. Solid bars = Realistic Model 3; hatched bars = Simplified Model 2. Error bars indicate the standard deviation across individual cells. Percentage labels denote the relative difference in mean dose for the realistic model compared with the simplified model.*

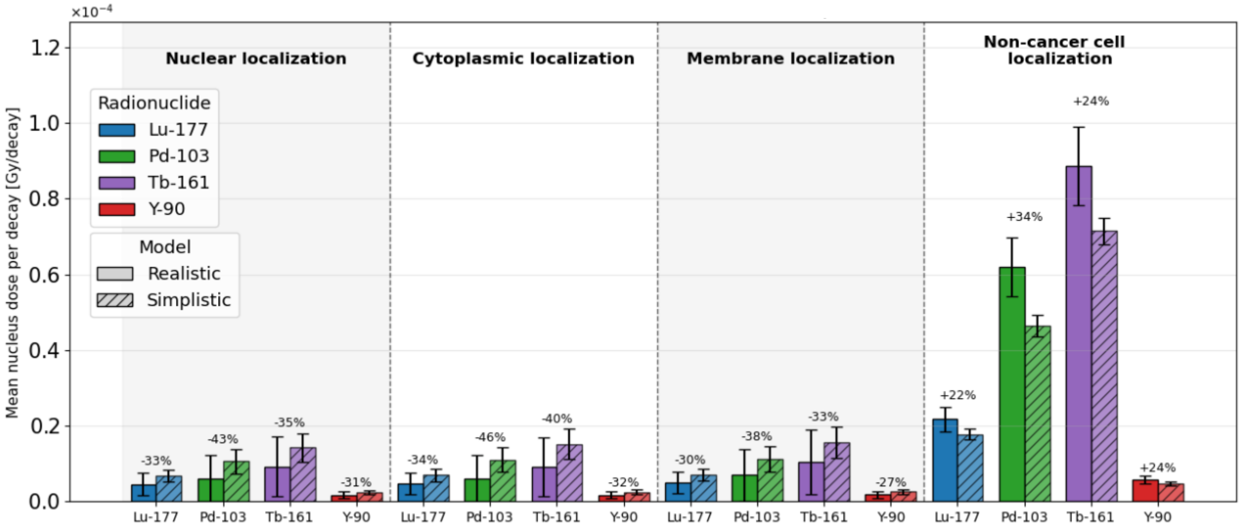


***Fig. S5.*** *Mean non-cancer-cell nuclear dose per cluster decay comparing Simplified Model 2 and Realistic Model 3. Solid bars = Realistic Model 3; hatched bars = Simplified Model 2. Error bars indicate the standard deviation across individual cells. Percentage labels denote the relative difference in mean dose for the realistic model compared with the simplified model.*

### Supplementary Table S6

##### Table S6a. Mean cancer-cell survival fractions for Simplified Model 2 and Realistic Model 2 at an initial cluster activity of 0.08 Bq.

| Radionuclide | Compartment | Mean SF (Simp.) | Mean SF (Real.) | SD SF (Simp.) | SD SF (Real.) | Diff. (%) |
|---|---|---|---|---|---|---|
| Lu-177 | Nucleus | 0.5855 | 0.5364 | 0.0318 | 0.0594 | -8.4% |
| Pd-103 | Nucleus | $1.13 \times 10^{-3}$ | $2.07 \times 10^{-4}$ | $8.73 \times 10^{-4}$ | $5.18 \times 10^{-4}$ | -81.7% |
| Tb-161 | Nucleus | 0.0318 | 0.0147 | $8.27 \times 10^{-3}$ | 0.0146 | -53.6% |
| Y-90 | Nucleus | 0.9530 | 0.9480 | $5.33 \times 10^{-3}$ | $9.28 \times 10^{-3}$ | -0.5% |
| Lu-177 | Cytoplasm | 0.7451 | 0.7854 | 0.0364 | 0.0538 | +5.4% |
| Pd-103 | Cytoplasm | 0.1606 | 0.2651 | 0.0689 | 0.1869 | +65.1% |
| Tb-161 | Cytoplasm | 0.3570 | 0.4848 | 0.0615 | 0.1406 | +35.8% |
| Y-90 | Cytoplasm | 0.9654 | 0.9697 | $5.46 \times 10^{-3}$ | $7.24 \times 10^{-3}$ | +0.4% |
| Lu-177 | Membrane | 0.7711 | 0.7913 | 0.0382 | 0.0468 | +2.6% |
| Pd-103 | Membrane | 0.2075 | 0.2684 | 0.0857 | 0.1369 | +29.3% |
| Tb-161 | Membrane | 0.4358 | 0.4969 | 0.0715 | 0.1121 | +14.0% |
| Y-90 | Membrane | 0.9676 | 0.9700 | $5.51 \times 10^{-3}$ | $6.55 \times 10^{-3}$ | +0.3% |
| Lu-177 | Non-cancer | 0.8359 | 0.8515 | 0.0395 | 0.0376 | +1.9% |
| Pd-103 | Non-cancer | 0.4384 | 0.4991 | 0.1499 | 0.1544 | +13.8% |
| Tb-161 | Non-cancer | 0.6593 | 0.6878 | 0.0894 | 0.0954 | +4.3% |
| Y-90 | Non-cancer | 0.9756 | 0.9777 | $5.61 \times 10^{-3}$ | $5.24 \times 10^{-3}$ | +0.2% |

*Simp. = Simplified Model 2; Real. = Realistic Model 2. SF = survival fraction calculated using the linear-quadratic model. SD = standard deviation across individual cells. Compartment indicates the cellular source localization. Diff. = percentage difference of the realistic relative to the simplified mean SF.*

##### Table S6b. Mean non-cancer-cell survival fractions for Simplified Model 2 and Realistic Model 2 at an initial cluster activity of 0.08 Bq.

| Radionuclide | Compartment | Mean SF (Simp.) | Mean SF (Real.) | SD SF (Simp.) | SD SF (Real.) | Diff. (%) |
|---|---|---|---|---|---|---|
| Lu-177 | Nucleus | 0.8661 | 0.8820 | 0.0313 | 0.0443 | +1.8% |
| Pd-103 | Nucleus | 0.5315 | 0.6044 | 0.1218 | 0.1978 | +13.7% |
| Tb-161 | Nucleus | 0.7220 | 0.7526 | 0.0698 | 0.1131 | +4.2% |
| Y-90 | Nucleus | 0.9800 | 0.9823 | $4.47 \times 10^{-3}$ | $6.17 \times 10^{-3}$ | +0.2% |
| Lu-177 | Cytoplasm | 0.8633 | 0.8700 | 0.0320 | 0.0533 | +0.8% |
| Pd-103 | Cytoplasm | 0.5160 | 0.5440 | 0.1237 | 0.2332 | +5.4% |
| Tb-161 | Cytoplasm | 0.7066 | 0.7005 | 0.0738 | 0.1647 | -0.9% |
| Y-90 | Cytoplasm | 0.9797 | 0.9813 | $4.54 \times 10^{-3}$ | $6.78 \times 10^{-3}$ | +0.2% |
| Lu-177 | Membrane | 0.8620 | 0.8615 | 0.0321 | 0.0562 | -0.1% |
| Pd-103 | Membrane | 0.5092 | 0.4968 | 0.1251 | 0.2367 | -2.4% |
| Tb-161 | Membrane | 0.7003 | 0.6668 | 0.0758 | 0.1690 | -4.8% |
| Y-90 | Membrane | 0.9795 | 0.9802 | $4.58 \times 10^{-3}$ | $7.15 \times 10^{-3}$ | +0.1% |
| Lu-177 | Non-cancer | 0.6852 | 0.6609 | 0.0236 | 0.0418 | -3.6% |
| Pd-103 | Non-cancer | 0.0205 | $8.27 \times 10^{-3}$ | $7.24 \times 10^{-3}$ | $8.16 \times 10^{-3}$ | -59.6% |
| Tb-161 | Non-cancer | 0.1273 | 0.0943 | 0.0172 | 0.0367 | -26.0% |
| Y-90 | Non-cancer | 0.9639 | 0.9612 | $3.74 \times 10^{-3}$ | $6.05 \times 10^{-3}$ | -0.3% |

*Simp. = Simplified Model 2; Real. = Realistic Model 2. SF = survival fraction calculated using the linear-quadratic model. SD = standard deviation across individual cells. Compartment indicates the cellular source localization. Diff. = percentage difference of the realistic relative to the simplified mean SF.*

### Supplementary Table S7

##### Table S7a. Mean cancer-cell survival fractions for Simplified Model 2 and Realistic Model 3 at an initial cluster activity of 0.1 Bq.

| Radionuclide | Compartment | Mean SF (Simp.) | Mean SF (Real.) | SD SF (Simp.) | SD SF (Real.) | Diff. (%) |
|---|---|---|---|---|---|---|
| Lu-177 | Nucleus | 0.5132 | 0.5068 | 0.0354 | 0.0745 | -1.2% |
| Pd-103 | Nucleus | $1.42 \times 10^{-4}$ | $4.92 \times 10^{-4}$ | $1.58 \times 10^{-4}$ | $1.87 \times 10^{-3}$ | +246.5% |
| Tb-161 | Nucleus | 0.0116 | 0.0178 | $4.14 \times 10^{-3}$ | 0.0201 | +53.6% |
| Y-90 | Nucleus | 0.9431 | 0.9393 | $6.41 \times 10^{-3}$ | 0.0126 | -0.4% |
| Lu-177 | Cytoplasm | 0.6910 | 0.7020 | 0.0430 | 0.1217 | +1.6% |
| Pd-103 | Cytoplasm | 0.0934 | 0.2222 | 0.0550 | 0.2558 | +137.9% |
| Tb-161 | Cytoplasm | 0.2681 | 0.3762 | 0.0606 | 0.2584 | +40.3% |
| Y-90 | Cytoplasm | 0.9574 | 0.9571 | $6.41 \times 10^{-3}$ | 0.0169 | -0.0% |
| Lu-177 | Membrane | 0.7199 | 0.7016 | 0.0440 | 0.1124 | -2.5% |
| Pd-103 | Membrane | 0.1292 | 0.2035 | 0.0726 | 0.2424 | +57.5% |
| Tb-161 | Membrane | 0.3448 | 0.3587 | 0.0748 | 0.2466 | +4.0% |
| Y-90 | Membrane | 0.9601 | 0.9571 | $6.54 \times 10^{-3}$ | 0.0154 | -0.3% |
| Lu-177 | Non-cancer | 0.7971 | 0.8429 | 0.0502 | 0.0820 | +5.7% |
| Pd-103 | Non-cancer | 0.3486 | 0.5336 | 0.1642 | 0.2751 | +53.1% |
| Tb-161 | Non-cancer | 0.5871 | 0.6807 | 0.1086 | 0.1912 | +15.9% |
| Y-90 | Non-cancer | 0.9697 | 0.9761 | $7.25 \times 10^{-3}$ | 0.0115 | +0.7% |

*Simp. = Simplified Model 2; Real. = Realistic Model 3. SF = survival fraction calculated using the linear-quadratic model. SD = standard deviation across individual cells. Compartment indicates the cellular source localization. Diff. = percentage difference of the realistic relative to the simplified mean SF.*

##### Table S7b. Mean non-cancer-cell survival fractions for Simplified Model 2 and Realistic Model 3 at an initial cluster activity of 0.1 Bq.

| Radionuclide | Compartment | Mean SF (Simp.) | Mean SF (Real.) | SD SF (Simp.) | SD SF (Real.) | Diff. (%) |
|---|---|---|---|---|---|---|
| Lu-177 | Nucleus | 0.8353 | 0.8899 | 0.0368 | 0.0695 | +6.5% |
| Pd-103 | Nucleus | 0.4497 | 0.6898 | 0.1295 | 0.2651 | +53.4% |
| Tb-161 | Nucleus | 0.6630 | 0.7839 | 0.0798 | 0.1659 | +18.2% |
| Y-90 | Nucleus | 0.9753 | 0.9829 | $5.38 \times 10^{-3}$ | $9.88 \times 10^{-3}$ | +0.8% |
| Lu-177 | Cytoplasm | 0.8320 | 0.8883 | 0.0380 | 0.0669 | +6.8% |
| Pd-103 | Cytoplasm | 0.4341 | 0.6988 | 0.1320 | 0.2606 | +61.0% |
| Tb-161 | Cytoplasm | 0.6457 | 0.7867 | 0.0844 | 0.1613 | +21.8% |
| Y-90 | Cytoplasm | 0.9748 | 0.9828 | $5.57 \times 10^{-3}$ | $9.26 \times 10^{-3}$ | +0.8% |
| Lu-177 | Membrane | 0.8302 | 0.8802 | 0.0379 | 0.0658 | +6.0% |
| Pd-103 | Membrane | 0.4272 | 0.6545 | 0.1324 | 0.2672 | +53.2% |
| Tb-161 | Membrane | 0.6382 | 0.7589 | 0.0858 | 0.1708 | +18.9% |
| Y-90 | Membrane | 0.9747 | 0.9816 | $5.55 \times 10^{-3}$ | $9.07 \times 10^{-3}$ | +0.7% |
| Lu-177 | Non-cancer | 0.6031 | 0.5304 | 0.0283 | 0.0585 | -12.0% |
| Pd-103 | Non-cancer | $3.33 \times 10^{-3}$ | $3.91 \times 10^{-4}$ | $1.88 \times 10^{-3}$ | $6.89 \times 10^{-4}$ | -88.3% |
| Tb-161 | Non-cancer | 0.0517 | 0.0216 | 0.0104 | 0.0144 | -58.2% |
| Y-90 | Non-cancer | 0.9536 | 0.9427 | $4.80 \times 10^{-3}$ | 0.0101 | -1.1% |

*Simp. = Simplified Model 2; Real. = Realistic Model 3. SF = survival fraction calculated using the linear-quadratic model. SD = standard deviation across individual cells. Compartment indicates the cellular source localization). Diff. = percentage difference of the realistic relative to the simplified mean SF.*

### Supplementary Table S8

**Table S8. Mean cancer-cell nuclear absorbed dose per total cluster decay for Simplified Model 3 and Realistic Model 3, used to characterize intra-voxel dose heterogeneity.**

| Radionuclide | Source localization | Mean dose (Real.) [Gy decay$^{-1}$] | Mean dose (Simp.) [Gy decay$^{-1}$] | Relative difference (%) |
|---|---|---|---|---|
| Lu-177 | Nucleus | $1.80\times10^{-5}$ | $1.22\times10^{-5}$ | +47.6 |
| Pd-103 | Nucleus | $4.86\times10^{-5}$ | $2.83\times10^{-5}$ | +71.5 |
| Tb-161 | Nucleus | $6.98\times10^{-5}$ | $4.26\times10^{-5}$ | +63.9 |
| Y-90 | Nucleus | $4.95\times10^{-6}$ | $3.55\times10^{-6}$ | +39.3 |
| Lu-177 | Cytoplasm | $1.03\times10^{-5}$ | $8.75\times10^{-6}$ | +18.2 |
| Pd-103 | Cytoplasm | $1.92\times10^{-5}$ | $1.51\times10^{-5}$ | +26.7 |
| Tb-161 | Cytoplasm | $2.72\times10^{-5}$ | $2.23\times10^{-5}$ | +22.0 |
| Y-90 | Cytoplasm | $3.50\times10^{-6}$ | $2.99\times10^{-6}$ | +17.1 |
| Lu-177 | Membrane | $1.03\times10^{-5}$ | $8.24\times10^{-6}$ | +25.2 |
| Pd-103 | Membrane | $1.93\times10^{-5}$ | $1.43\times10^{-5}$ | +34.9 |
| Tb-161 | Membrane | $2.77\times10^{-5}$ | $2.02\times10^{-5}$ | +36.8 |
| Y-90 | Membrane | $3.49\times10^{-6}$ | $2.88\times10^{-6}$ | +21.1 |
| Lu-177 | Non-cancer | $5.26\times10^{-6}$ | $1.22\times10^{-5}$ | -56.9 |
| Pd-103 | Non-cancer | $7.66\times10^{-6}$ | $2.83\times10^{-5}$ | -72.9 |
| Tb-161 | Non-cancer | $1.11\times10^{-5}$ | $4.26\times10^{-5}$ | -73.9 |
| *Y-90* | Non-cancer | *$1.95\times10^{-6}$* | $3.55\times10^{-6}$ | -45.2 |

*Note: Relative difference is calculated as (Realistic − Simplified) / Simplified × 100. Non-cancer localization values are compared with nuclear localization of the respective radionuclide.*

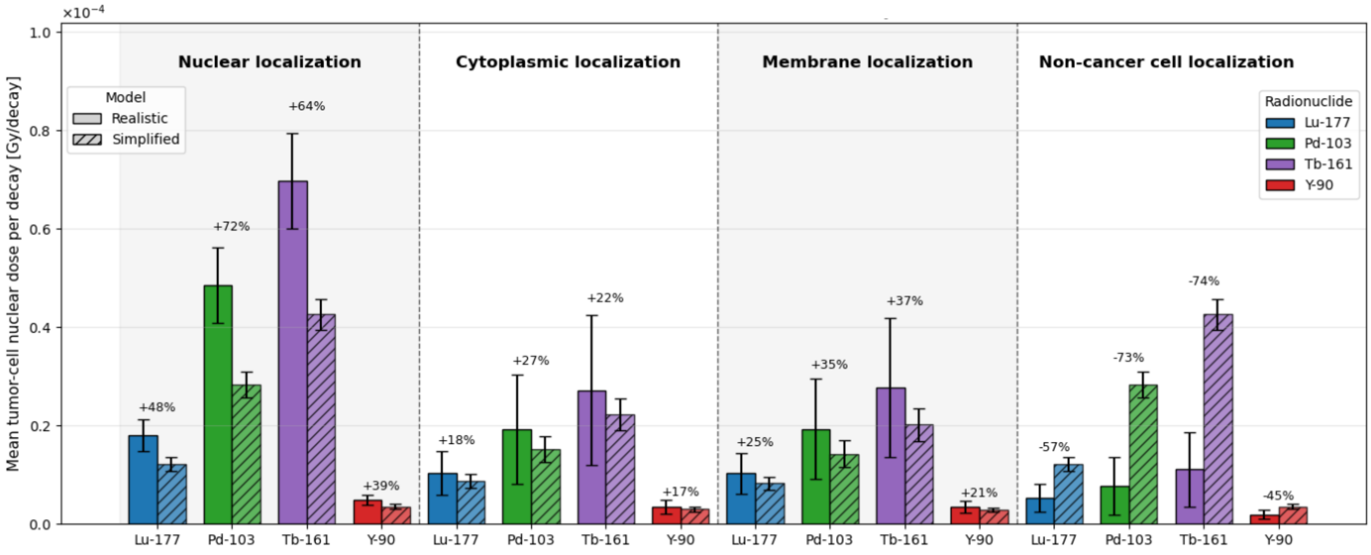


***Fig. S6.*** *Mean cellular nuclear dose per cluster decay comparing Simplified Model 3 and Realistic Model 3 used to examine intra-voxel heterogeneity. Solid bars = Realistic Model 3; hatched bars = Simplified Model 3. Error bars indicate the standard deviation across individual cells. Percentage labels denote the relative difference in mean dose for the realistic model compared with the simplified model.*

#### Supplementary Table S9

**Table S9. Physical decay characteristics of the radionuclides used in this study. *Values mentioned are the weighted averages in keV per decay, unless stated otherwise. Adapted from the supplementary material of Fourie et al. [5].**

| Property | Lu-177 | Y-90 | Pd-103 (+Rh-103m) | Tb-161 |
|---|---|---|---|---|
| **Half-life** | 6.647 d | 2.671 d | 16.991 d | 6.95 d |
| **Principal decay mode** | β− decay | β− decay | EC & isomeric transition | β− decay |
| **Daughter** | Hf-177 (stable) | Zr-90 (stable) | Rh-103m → Rh-103 (stable) | Dy-161 (stable) |
| **Mean Auger-electron (AE) energy** | 1.13 | 0.0007 | 8.54 | 8.94 |
| **Number of AE per decay** | 1.2 | — | 13.3 | 11.0 |
| **AE energy range (mm)** | 0.01–61.7 | — | 0.034–22.3 | 0.018–50.9 |
| **Mean conversion-electron (CE) energy** | 13.52 | 0.2 | 34.97 | 39.28 |
| **CE energy range (mm)** | 6.2–206 | — | 16.6–39.8 | 3.3–98.3 |
| **Mean β energy** | 133.3 | 932.9 | — | 154.3 |
| **Total electron energy** | 147.9 | 933.1 | 43.5 | 202.5 |
| **Total photon energy (X & γ)** | 35.1 | 0.0012 | 16.14 | 36.35 |
| **Total energy (photon + electron)** | 183 | 933.1 | 59.65 | 238.9 |
| **Photon/electron ratio** | 0.24 | ~0 | 0.37 | 0.18 |

*AE, Auger electron; CE, conversion electron; EC, electron capture.*
